# Provenance Attestation of Human Cells Using Physical Unclonable Functions

**DOI:** 10.1101/2021.06.11.448108

**Authors:** Yi Li, Mohammad Mahdi Bidmeshki, Taek Kang, Chance M. Nowak, Yiorgos Makris, Leonidas Bleris

## Abstract

We introduce a novel methodology, namely CRISPR-Engineered Attestation of Mammalian Cells using Physical Unclonable Functions (CREAM-PUFs), which can serve as the cornerstone for formally verifying transactions in human cell line distribution networks. A PUF is a physical entity which provides a measurable output that can be used as a unique and irreproducible identifier for the artifact wherein it is embedded. Popularized by the electronics industry, silicon PUFs leverage the inherent physical variations of semiconductor manufacturing to establish intrinsic security primitives for attesting integrated circuits. Owing to the stochastic nature of these variations and the multitude of steps involved, photo-lithographically manufactured silicon PUFs are impossible to reproduce (thus unclonable). Inspired by the success of silicon PUFs, we sought to exploit a combination of sequence-restricted barcodes and the inherent stochasticity of CRISPR-induced non-homologous end joining DNA error repair to create the first generation of genetic physical unclonable functions in three distinct human cells (HEK293, HCT116, and HeLa). We demonstrate that these CREAM-PUFs are robust (i.e., they repeatedly produce the same output), unique (i.e., they do not coincide with any other identically produced PUF), and unclonable (i.e., they are virtually impossible to replicate). Accordingly, CREAM-PUFs can serve as a foundational principle for establishing provenance attestation protocols for protecting intellectual property and confirming authenticity of engineered cell lines.

## Introduction

Recent advances in synthetic biology and genome editing^1–9^ have enabled development of a broad range of engineered cells and have fueled emergence of a novel industry which seeks to produce specialized cell lines^10–13^ and monetize them through commercial distribution networks. Many such highly customized proprietary cell lines are the result of extensive and expensive research and development efforts and come with price tags in the tens of thousands of dollars. Therefore, the legitimate producers of these valuable cell lines have a vested interest to protect their intellectual property and recover their investment by ensuring that their proprietary cell line does not get illicitly copied and distributed. At the same time, customers who acquire such expensive cell lines also have a vested interest in being assured of the origin (and, thereby, the quality) of their purchase, as well as holding proof of legitimate ownership of the cell line. In short, this emerging industry is in need of novel protocols for formally verifying the sale transaction of proprietary cell lines.

Moreover, cross-contamination or misidentification of cell lines due to poor handling, mislabeling, or procurement from dubious or undocumented sources is a rampant problem, resulting in innumerable financial and time losses^14–19^. For example, a major German cell repository has reported that 20% of its human cell line stocks were cross contaminated with other cell lines, and the China Center for Type Culture Collection demonstrated that 85% of cell lines in their repository, supposedly established from primary isolates, were actually HeLa cells. Such issues undermine quality, repeatability and, ultimately, overall efficiency of medical research. Therefore, quality control and source verification provisions are paramount toward safeguarding against working with unsuitable cell line models and producing false data.

To this end, we introduce CRISPR Engineered Authentication of Mammalian Cells (CREAM-PUFs), a methodology which enables *provenance attestation* of human cell lines through the use of the first genetic Physical Unclonable Functions (PUFs). A PUF is a hardware security primitive which exploits the inherent randomness of its manufacturing process to enable attestation of the entity wherein it is embodied^20–23^. A PUF is typically modeled as a mapping between input stimuli (challenges) and output values (responses), which is established stochastically among a vast array of options and is, therefore, unique and irreproducible. Upon manufacturing, a PUF is interrogated and a database comprising valid Challenge-Response Pairs (CRPs) produced by this PUF is populated (**Figure 1**). Attestation can, thus, be achieved by issuing a challenge to the holder of the physical entity embodying the PUF, receiving the response and comparing against the golden references stored in the database. Accordingly, typical quality metrics for evaluating a PUF include *robustness,* i.e., the probability that given the same challenge it will consistently produce the same response, and *uniqueness,* i.e., the probability that its mapping does not coincide with the mapping of any other identically manufactured PUF.

**Fig. 1.**
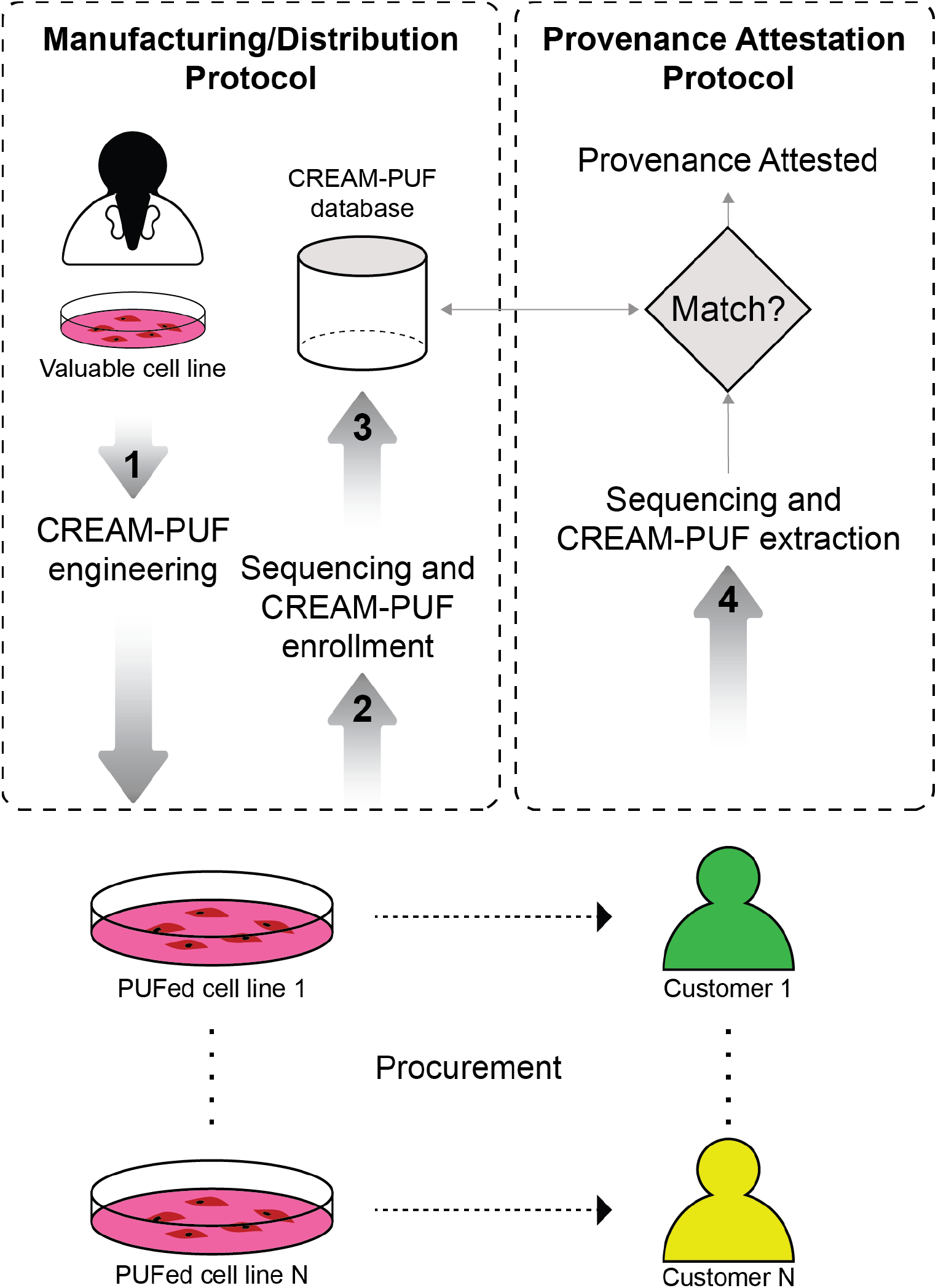
Provenance attestation protocols and pilot CREAM-PUF. **(A)** The producer of a valuable cell line inserts a unique, robust and unclonable signature in each legitimately produced copy of this cell line. Upon thawing of a frozen sample and prior to its initial use, a customer who purchased a copy of the cell line can obtain this signature and communicate it to the producer who compares it against the signature database of legitimately produced copies of this cell line and, thereby, attests its provenance.

While PUF-like concepts were proposed earlier in the literature, their popularity soared after their first implementation in silicon, as part of electronic integrated circuits^24^. Indeed, by exploiting the inherent variation of advanced semiconductor manufacturing processes, silicon PUFs became a commercial success, serving as the foundation of many security protocols implemented both in software and in hardware. While this success stimulated similar efforts in various other domains, to date PUFs have yet to be adopted in the context of biological sciences, wherein they could find numerous applications. Similar to the use of silicon PUFs (in their simplest form) as unique IDs for verifying genuineness of electronic circuits, genetic PUFs could be embedded in cell lines to attest their provenance.

More specifically, CREAM-PUFs could enable the producer of a valuable cell line to insert a unique, robust and unclonable signature in each legitimately produced copy of this cell line. Upon thawing of a frozen sample and prior to its initial use, a customer who purchased a copy of the cell line can obtain this signature and communicate it to the producer who compares it against the signature database of legitimately produced copies of this cell line and, thereby, attests its provenance (**Figure 1**). Through this protocol, the producer of the cell line can ensure that anyone publicly claiming ownership of a copy of this cell line has acquired it legitimately. At the same time, the customer can be assured of the source and quality of the procured cell line, as the producer explicitly confirms its origin and assumes responsibility for its production.

Toward developing CREAM-PUFs, we hypothesized that a process which combines molecular barcoding with non-homologous end joining (NHEJ) repair and exploits the inherent stochasticity of the latter **(Figure 2A)**, yields measurable genetic changes that satisfy all PUF conditions. More specifically, a two-dimensional mapping between barcodes and indels resulting from this process, which can be obtained by sequencing a genetic locus of the cell line, is a *robust* yet *unique* and *unclonable* signature.

**Fig. 2.**
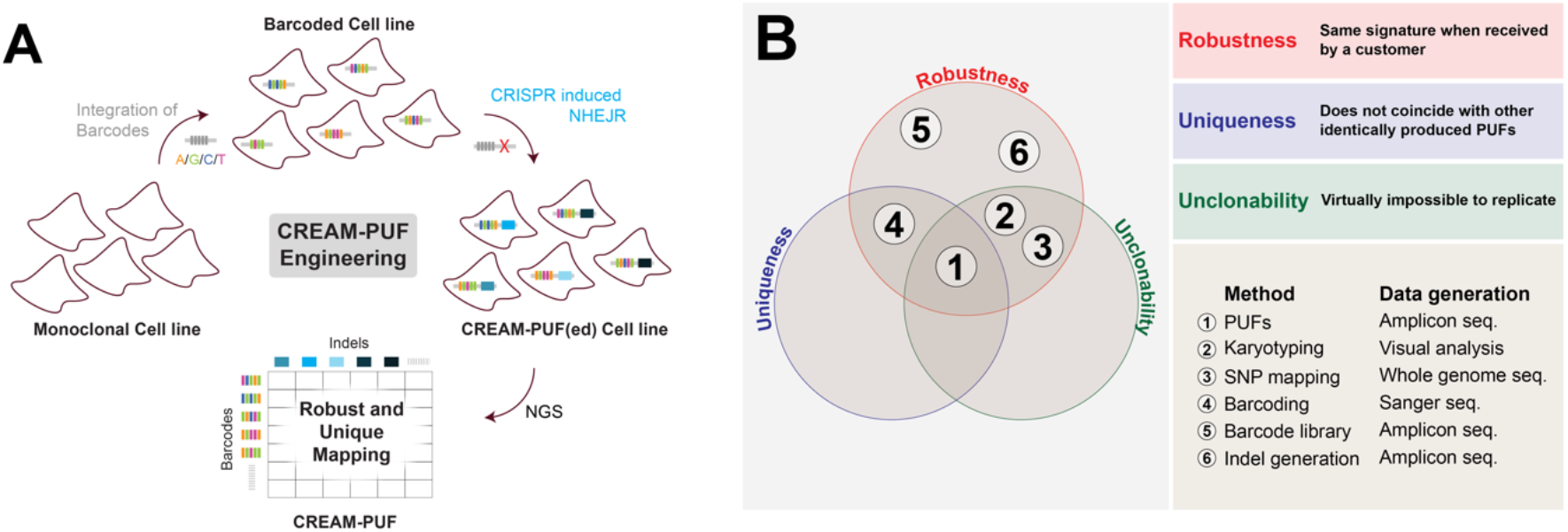
Overview of the CREAM-PUF generation process. **(A)** Schematic illustration of design of CREAM-PUFs. Barcodes were stably integrated into cell lines of interest, which were subsequently subjected to CRISPR/SpCas9 treatment to induce non-homologous end joining (NHEJ). The resulting two-dimensional mapping between barcodes and indels is evaluated for robustness and uniqueness. **(B)** Venn diagram comparing PUFs to other methods/technologies.

As visualized in a Venn diagram (**Figure 2B**), CREAM-PUFs is the only methodology that satisfies all three PUF criteria. Barcodes and indels alone are not PUFs and cannot be used for provenance attestation. Indels are not PUFs because they are not unique^25,26^ and are clonable (thus violate two of the three PUF conditions). Barcodes are also not PUFs, as they violate the uniqueness criterion. Indeed, as shown later in this manuscript, when we integrated a 5-nucleotide barcode library into the *AAVS1* locus of human HEK293 cells via CRISPR/SpCas9 in six parallel replicates, we observed that the uniqueness criterion is not satisfied. We emphasize that increasing the size of the barcode would not resolve the uniqueness criterion but would merely increase complexity. In contract, the uniqueness of our PUF design is not based on a scalar property, such as the complexity or entropy of barcodes or indels, but rather on the joint probability distributions of both barcodes and indels in the cell population. Finally, natural genetic variations such as short nucleotide polymorphisms (SNPs) or short tandem repeats (STRs) can indeed be used for cell line authentication but not for provenance attestation, because they are, generally, not unique or unclonable (**Figure 2B**). As an example, all cell lines derived from a single monoclonal source share the same SNP mapping or karyotyping information and thus violate the uniqueness requirement.

## Results

To validate our hypothesis, and towards implementing the first generation of genetic PUFs, we carried out a pilot study where we leverage genome engineering using Clustered Regularly Interspaced Short Palindromic Repeats (CRISPR)^27–30^. CRISPR is an immune response mechanism against bacteriophage infections in bacteria and archaea that has revolutionized the field of genome editing and spurred myriads of applications critically relevant to agriculture, biomanufacturing, and human health^28,31–35^. Critically, Cas9 can be programmed to bind to a specific region of DNA and generate a double stranded break which, in turn, initiates the error prone DNA repair pathway NHEJ. Our method involved the following steps.

First, we stably integrated a 5-nucleotide barcode library into the *AAVS1* locus of human HEK293 cells via CRISPR/SpCas9-mediated homologous recombination (HR). Specifically, as shown in **Figure 3**, a 5-nucleotide barcode (5’-NNNNN-3’, complexity: 4^5^ = 1024) was placed immediately upstream of a truncated CMV (225 bp versus 612 bp of the full-length CMV promoter) mKate construct and a PGK1-hygromycin resistance gene for drug selection. The safe harbor *AAVS1* locus was chosen as the integration site to minimize potential disruption of normal cellular functions upon the stable integration of the transgenes^36^. Subsequently, the genomic DNA from the resulting stable cells was collected and used as the PCR template (**Supplementary Table 1**, primers P1 and P2) to isolate the cDNA transcript harboring the barcode, which were subsequently subjected to NGS (Next Generation Sequencing)-based amplicon sequencing. In total, 805 distinct barcodes were detected (**Figure 4 and Supplementary Table 2**).

**Fig 3.**
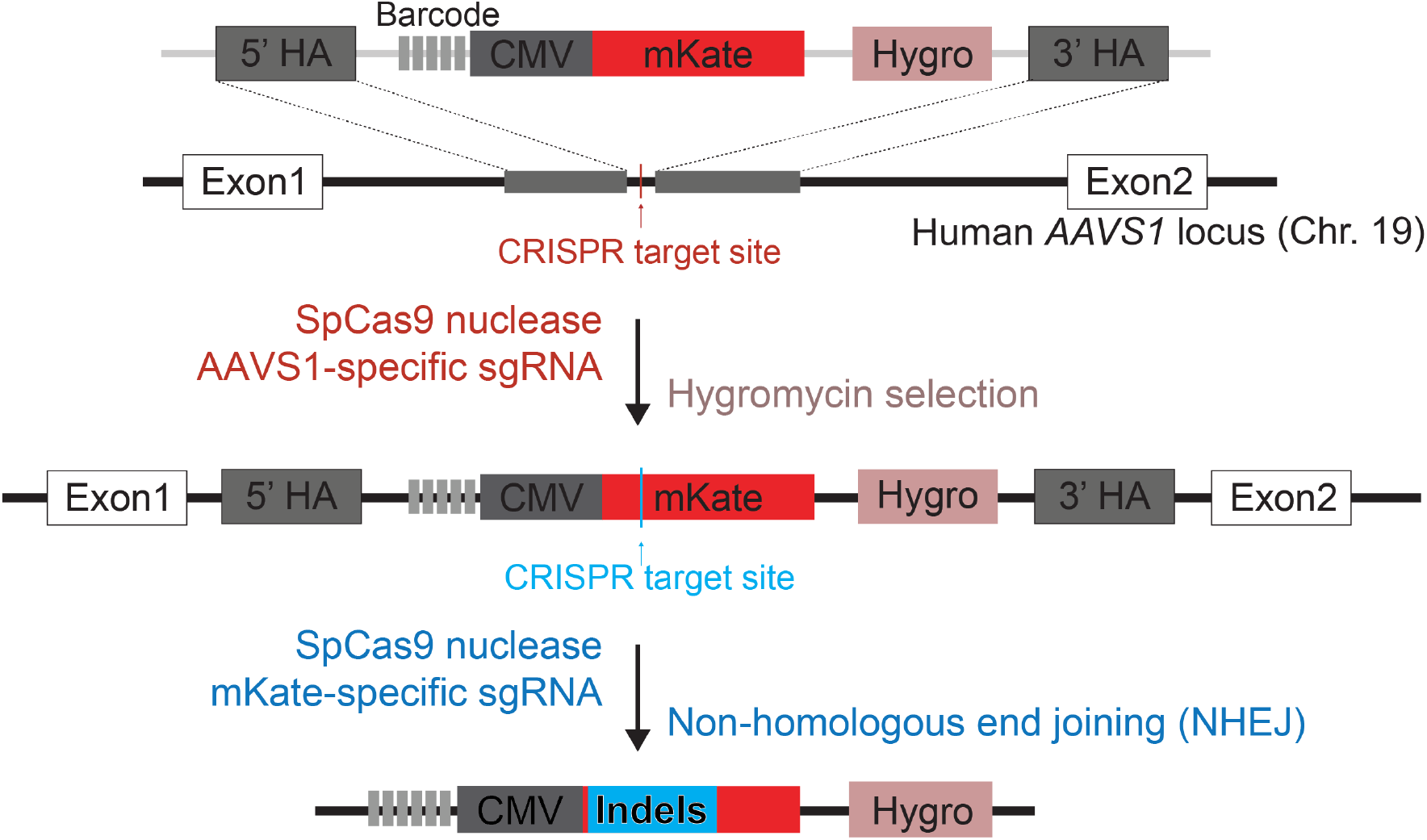
Schematic illustration of implementation of CREAM-PUFs. Using CRISPR-Cas9 system, a set of synthetic constructs containing an array of 5-bp barcodes (5’-NNNNN-3’), constitutive fluorescent reporter and hygromycin resistance gene were stably integrated into the human *AAVS1* safe harbor locus. Next, the cells were transiently transfected with CRISPR to induce NHEJ.

**Fig. 4.**
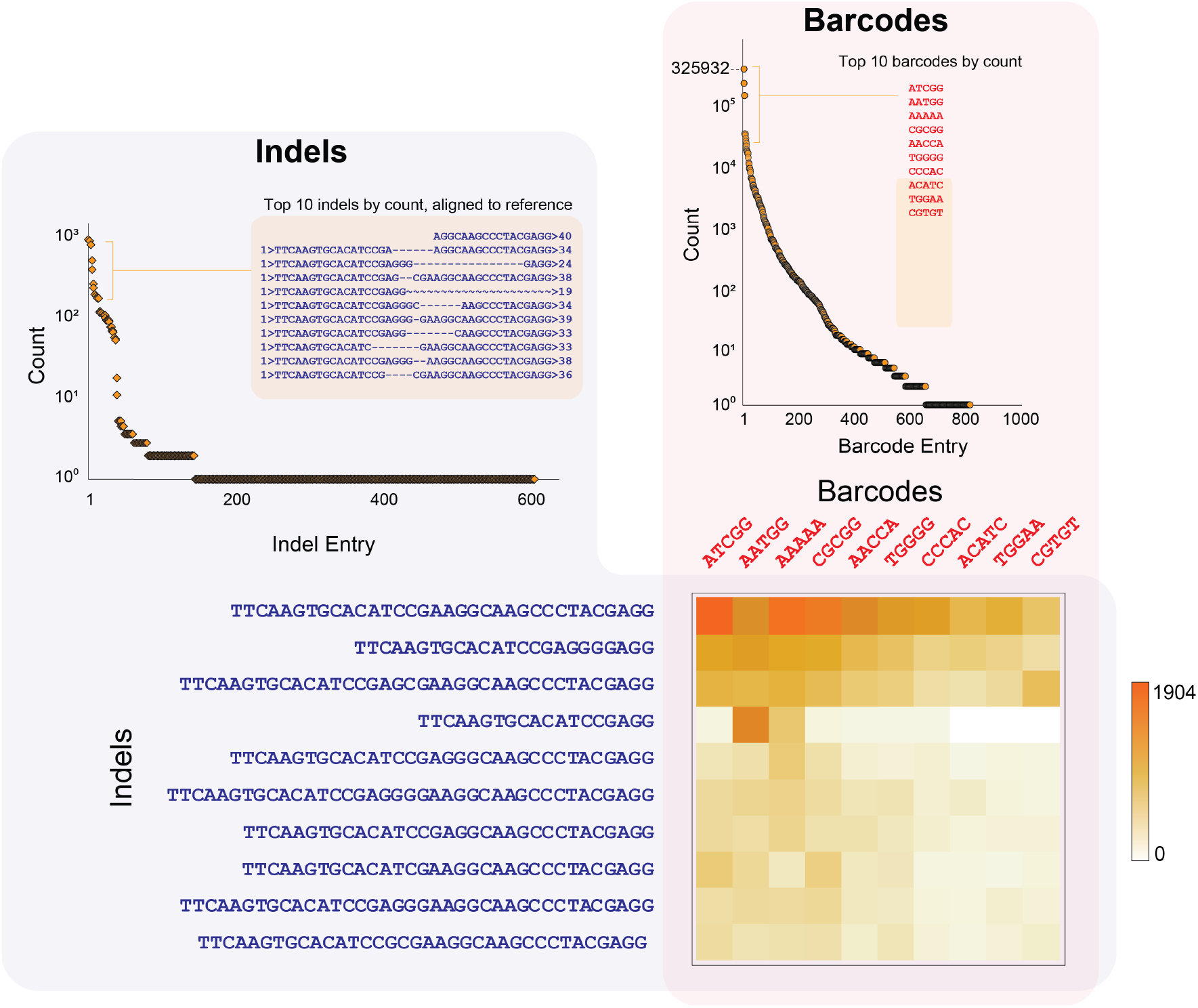
Distribution of indels and barcodes that makeup a CREAM-PUF (Right) Frequency of detected barcodes. In total, 805 unique barcodes were observed. **(Left)** Frequency of detected indels. In total, 569 unique indels were observed. **(Bottom)** Barcode/Indel matrix. Heatmap presentation of the pilot CREAM-PUF matrix consisting of the 10 most frequently occurring barcodes and indels.

Next, we aimed to combine the randomness of transfection into the barcoded cells and the inherent stochasticity of the cellular DNA error-repair processes to create a unique two-dimensional mapping between the barcodes and the indels. To this end, we initially screened five sgRNAs (**Supplementary Figure 1**)^37^ for targeting efficiency by designing the sgRNA to target the ORF of the fluorescence reporter mKate. As shown in **Supplementary Figure 1**, when co-transfected with SpCas9, all 5 sgRNAs efficiently suppressed the expression of mKate (sgRNA-5 was used for all subsequent experiments). We, therefore, proceeded by transiently transfecting the barcoded cell line with a sgRNA that targets adjacent to the integrated barcode in order to induce NHEJ repair.

Subsequently, the genomic DNA from the CRISPR-treated barcoded cell line was extracted and the amplicons containing both the barcodes and the expected indel sequences were prepared using PCR (primers P1 and P2). This was followed by NGS sequencing (100bp paired-end reads), which provided both the barcode sequence (forward end) and the indel sequence (reverse end). As shown in **Figure 4**, in total 569 distinct indels were observed (**Supplementary Table 3**) and the most frequently occurring indels demonstrated deletions of 1- to 16-nucleotides flanking the predicted SpCas9 cutting site (cutting site: between 5’-CGAGGG-3’ and 5’-CGAAGG-3’, PAM: AGG)^38^.

The detected indels were associated with their corresponding barcodes from the same reads and the resulting two-dimensional matrix was sorted by the frequencies of barcoded indels (**Supplementary Table 4**). As expected, CRISPR-mediated editing occurred in a subpopulation of a non-uniformly distributed barcoded cell population (**Supplementary Table 2**), resulting in 218 out of the total 805 barcodes being present in the barcode and indels matrix. We provide the cropped matrix for the most frequently detected barcode and indel sequences in **Figure 4.** By simple inspection, the utility of this matrix as a PUF to support CRispr-Engineered Authentication of Mammalian Cells (CREAM-PUF) becomes apparent: using silicon PUF terminology, a vector of (barcode, indel) elements in this matrix can be used as a *challenge,* while the corresponding vector of frequencies can be used as the *response*.

However, before relying on CREAM-PUFs for attesting provenance of a cell line, we sought to evaluate their aptitude as PUFs (**Figure 2B**). To this end, building upon our experience with the initial pilot experiments, we thoroughly assessed CREAM-PUFs using the strategy illustrated in **Figure 5.** With numerous PUFs constructed across various human cell lines, we performed individual PUF and pairwise comparisons to establish their *robustness* (i.e., their ability to produce matching signatures when a cell line is sequenced multiple times, e.g., at the vendor and at the customer site) and *uniqueness* (i.e., their ability to produce distinct signatures when multiple, identically produced copies of the same cell line are sequenced).

**Fig. 5.**
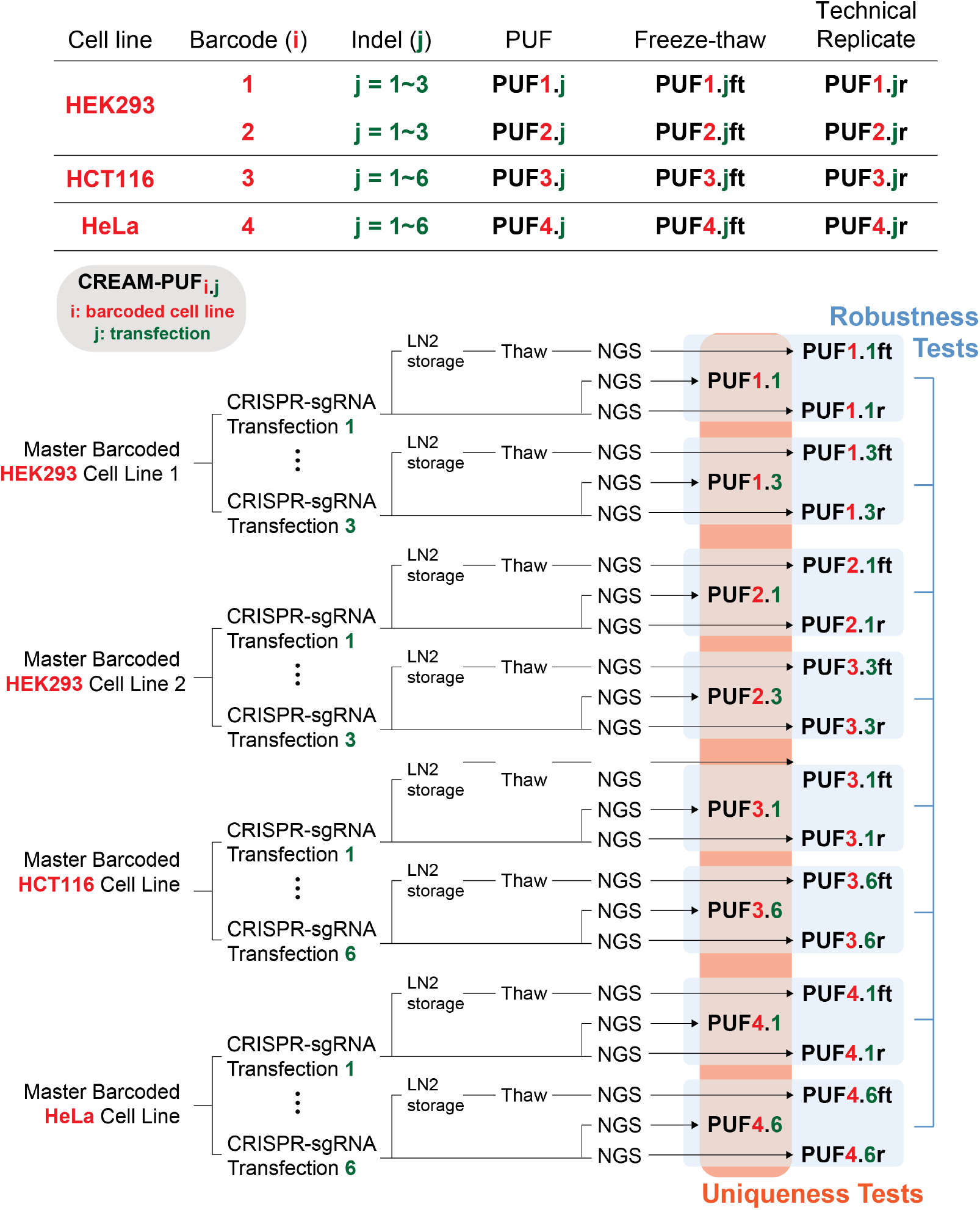
List of all CREAM-PUFs generated for this study. In this study, 2 independently barcoded HEK293 cell lines were each transfected with identical sgRNA in 3 separate instances, resulting in a total of 6 unique PUFs. In addition, a HCT116 cell line and a HeLa cell line, each with a distinct set of barcodes, were each transfected 6 times to generated 6 unique PUFs for each cell line. A portion of each PUF was subjected to one cycle of freeze-thaw (denoted as ft) before proceeding with next generation sequencing (NGS). In order to account for sequencing errors, PUFs from each barcoded cell line were also sequenced twice (denoted as r). Therefore, in total, 50 samples were sequenced via NGS, each resulting in a unique matrix of barcode/indel frequencies. To assess robustness to NGS measurement error, the matrix for each PUF i.j is compared against the matrix of its technical replicate PUF i.jr (i={1,2,3,4}). Similarly, to assess robustness to the freeze-thaw process, the matrix for each PUF i.j is compared against the matrix of its freeze-thaw counterpart PUF i.jft (i ={1,2,3,4}). To assess uniqueness, the matrices of all PUF i.j (i={1,2,3,4}) are compared pairwise.

To facilitate such comparisons, two independently engineered, barcoded cell lines (Barcoded Cell Line #1 and Barcoded Cell Line #2) were prepared for HEK293 cells. In parallel, two additional barcoded cell lines were also generated for HCT116 (Barcoded Cell Line #3) and HeLa (Barcodes Cell Line #4) cells, respectively. Next, for each of the two cell lines derived from HEK293, we transfected the barcoded cells with the same sgRNA (**Supplementary Figure 1,** sgRNA-5) three times (independent experiments), producing a total of 6 CREAM-PUFs (PUF1.1, PUF1.2 and PUF1.3 from Barcoded Cell Line #1, and PUF2.1, PUF2.2 and PUF2.3 from Barcoded Cell Line #2). We also subjected all engineered cells to one cycle of freezing and thawing, resulting in PUF1.1ft, PUF1.2ft, PUF1.3ft for Barcoded Cell Line #1 and PUF2.1ft, PUF2.2ft, PUF2.3ft for Barcoded Cell Line #2, respectively. These CREAM-PUFs were subjected to NGS analysis to produce the previously described barcode-indel matrix for each one of them. To incorporate and account for measurement errors introduced at the NGS step, PUF1.1 and PUF2.1 were sequenced twice, with the repeat results named PUF1.1r and PUF2.1r, respectively. Similarly, the two cell lines derived from HCT116 and HeLa were each subjected to 6 independent CRISPR/sgRNA treatments and the resulting cells (PUF3.j and PUF4.j, respectively) were subjected to one cycle of freezing and thawing (PUFi.jft, i={3,4}, j={1-6}), as well as repeated NGS sequencing (PUFi.jr, i={3,4}, j={1-6}). All the CREAM-PUFs produced by our experiments and used in our evaluation are summarized in **Figure 5**.

To evaluate robustness, we compare the NGS-generated barcode/indel matrix of PUFi.j to those of PUFi.jr and PUFi.jft (i={1,2,3,4}), anticipating that they match (**Figure 5, Robustness Tests**). Similarly, to evaluate CREAM-PUF uniqueness that stems from the stochastic nature of NHEJ repair and the random association with the barcodes, we compare the NGS-generated barcode/indel matrix across all PUFs (**Figure 5, Uniqueness Tests**) anticipating that they are distinct.

For a qualitative assessment, we focus on the most densely populated area of the barcode/indel matrix. As an example, in **Figure 6A** and **Figure 6B**, we provide the frequencies and sequences of the five most frequently observed barcodes and indels for PUFi.1, PUFi.1r and PUFi.1ft (i={1,2}, respectively) from HEK293 cells. We also provide heatmaps of the 30 most frequently observed barcodes and indels (complete sequence data in **Supplementary Tables 5-8**). These remain qualitatively the same and suggest a high level of robustness among these samples (which will be quantified in the following sections).

**Fig. 6.**
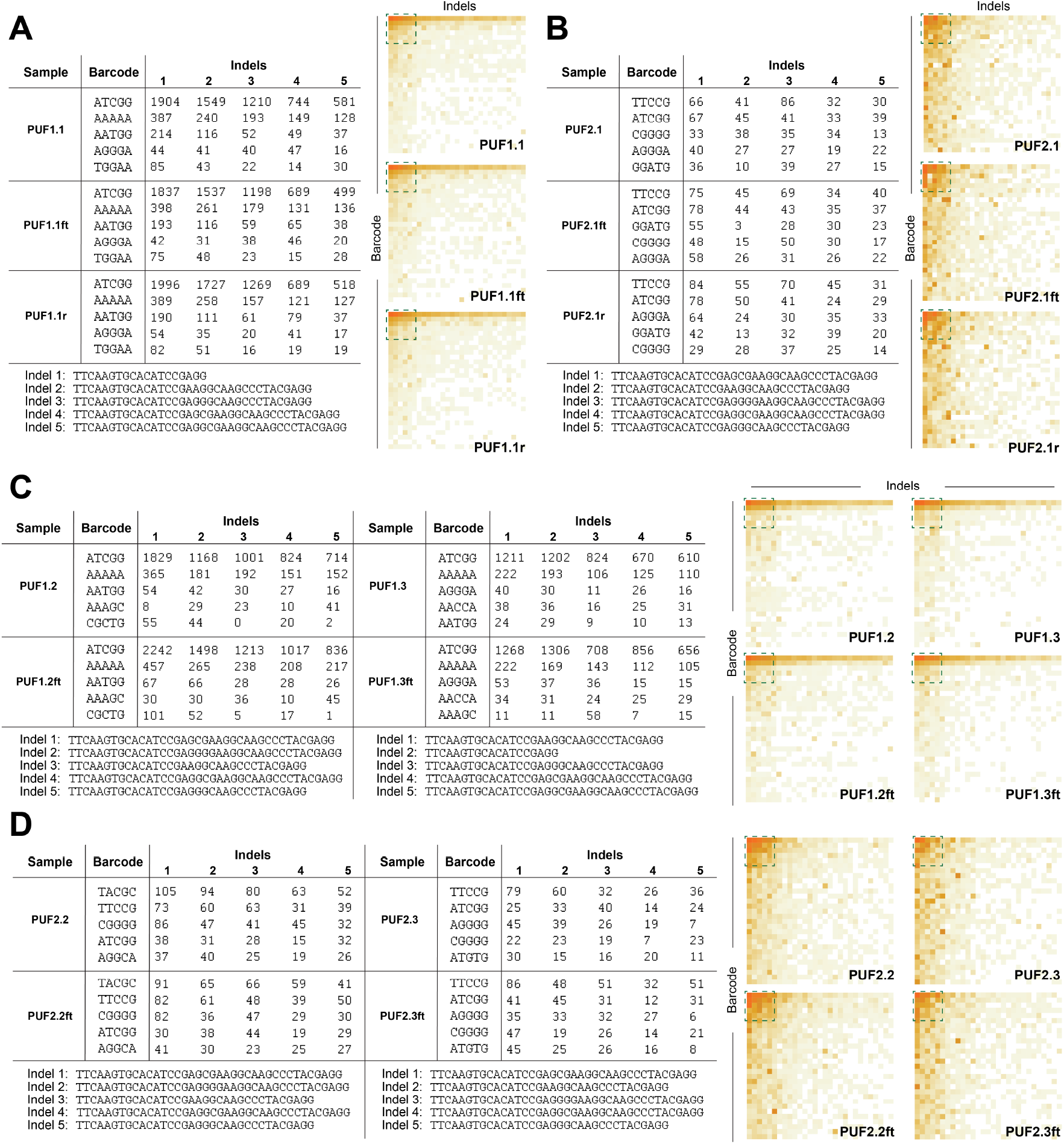
Qualitative assessment of CREAM-PUFs generated using HEK293. **(A~D)** Frequencies of barcode-indel addresses consisting of the 5 most commonly observed barcodes and indels (**Left**) and heatmap based on the same data but expanded to the top 30 most commonly observed barcodes and indels (**Right**) for a given PUF and its freeze-thaw counterparts and technical replicates (if applicable). The green dashed square on the heatmap represents the data shown on the table. Data shown in **(A, B)** are barcode-indel addresses for PUF1.1 **(A)** and PUF2.1 **(B)** with their respective freeze-thaw counterpart and technical replicate. Data shown in **(C, D)** are barcode-indel addresses for PUF1.2, PUF1.3 **(C)**, PUF2.2, PUF2.3 **(D)** with their respective freeze-thaw counterpart.

In contrast, different PUFs exhibit dissimilar patterns of the cropped CREAM-PUF matrices (e.g., PUF1.2 and PUF1.3 in **Figure 6C**) and, importantly, different representation in the most frequently observed barcodes and indels. As an example, the 3^rd^ and 4^th^ most frequently observed barcodes for PUF1.2 were 5’-AATGG-3’ and 5’-AAAGC-3’, while for PUF1.3 they were 5’-AGGGA-3’ and 5’-AACCA-3’, respectively. Similarly, the most frequent indel from PUF1.2 was 5’-TTCAAGTGCACATCCGAGG-3’, while for PUF1.3 it was 5’-TTCAAGTGCACATCCGAAGGCAAGCCCTACGAGG-3’ (**Supplementary Table 5**). These results suggest that a CREAM-PUF identifier based on a combination of the barcode/indel sequences and their respective counts can satisfy both robustness and uniqueness.

As mentioned earlier, we introduced 6 PUFs in each of two additional human cell lines (HCT116 and HeLa). Our sequencing results (all PUFs in **Supplementary Figures 2-3)** show that qualitatively these PUFs also satisfy both robustness and uniqueness. We provide the frequencies and sequences of the five most frequently observed barcodes and indels for representative PUFs for both cell lines (**Figure 7**). For example, for HCT116 cells, the heatmaps were visually similar among PUF3.2, PUF3.2ft and PUF3.2r, while being distinct between PUF3.2 and the rest of the PUFs **(Figure 7A, Supplementary Figure 2)**. Similarly, for HeLa cells, while the 5^th^ most frequently observed indels from PUF4.2, PUF4.2ft and PUF4.2r remained as 5’-CCTCGGATGTGCACTTGAA-3’, this sequence was not observed in the most frequent indel list (top 5) from PUF4.1 sample **(Figure 7B, Supplementary Figure 3)**. All barcode and indel sequences from HCT116 and HeLa are included in **Supplementary Tables 9-10**.

**Fig. 7.**
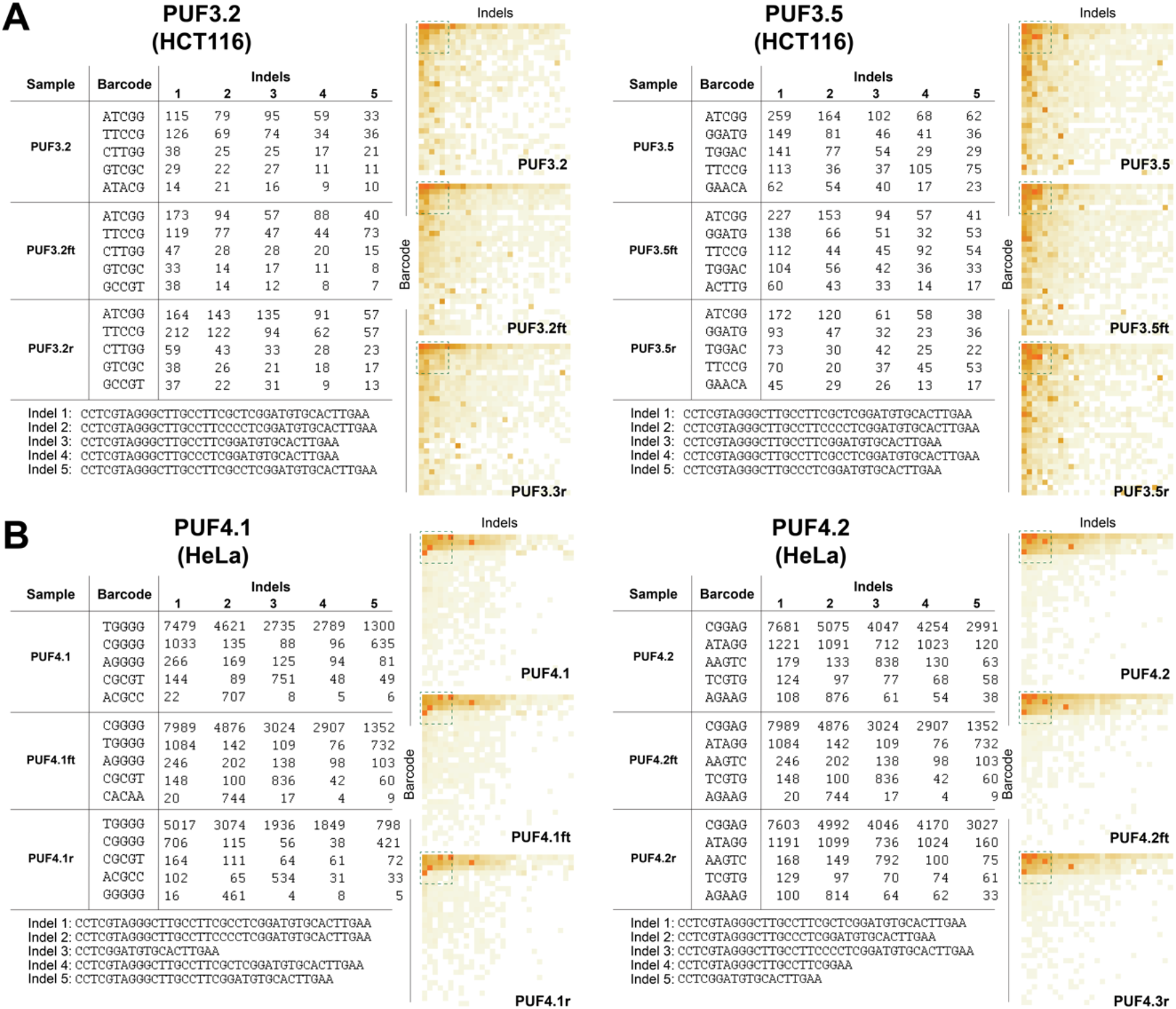
Qualitative assessment of CREAM-PUFs generated using HCT116 and HeLa. Qualitative analysis of PUFs as shown in **Figure 6**, with HCT116 (**A**) and HeLa (**B**). In both cell types, heatmaps of barcode-indel addresses from *intra*-PUFs were visually similar, via different between *inter*-PUFs.

For provenance attestation, the end-user of a CREAM-PUF(ed) cell line must provide the NGS data (i.e., barcode/indel matrix), which is then compared against the values stored in a database to determine whether there is a match. Importantly, to facilitate quantitative evaluation of the similarity between CREAM-PUF matrices, we first concatenate the barcode and indel sequences to generate unique addresses (**Figure 8).** This allows us to express each CREAM-PUF as a probability distribution (**Supplementary Table 11**), based on the frequency of occurrence for each unique barcode-indel address.

**Fig. 8.**
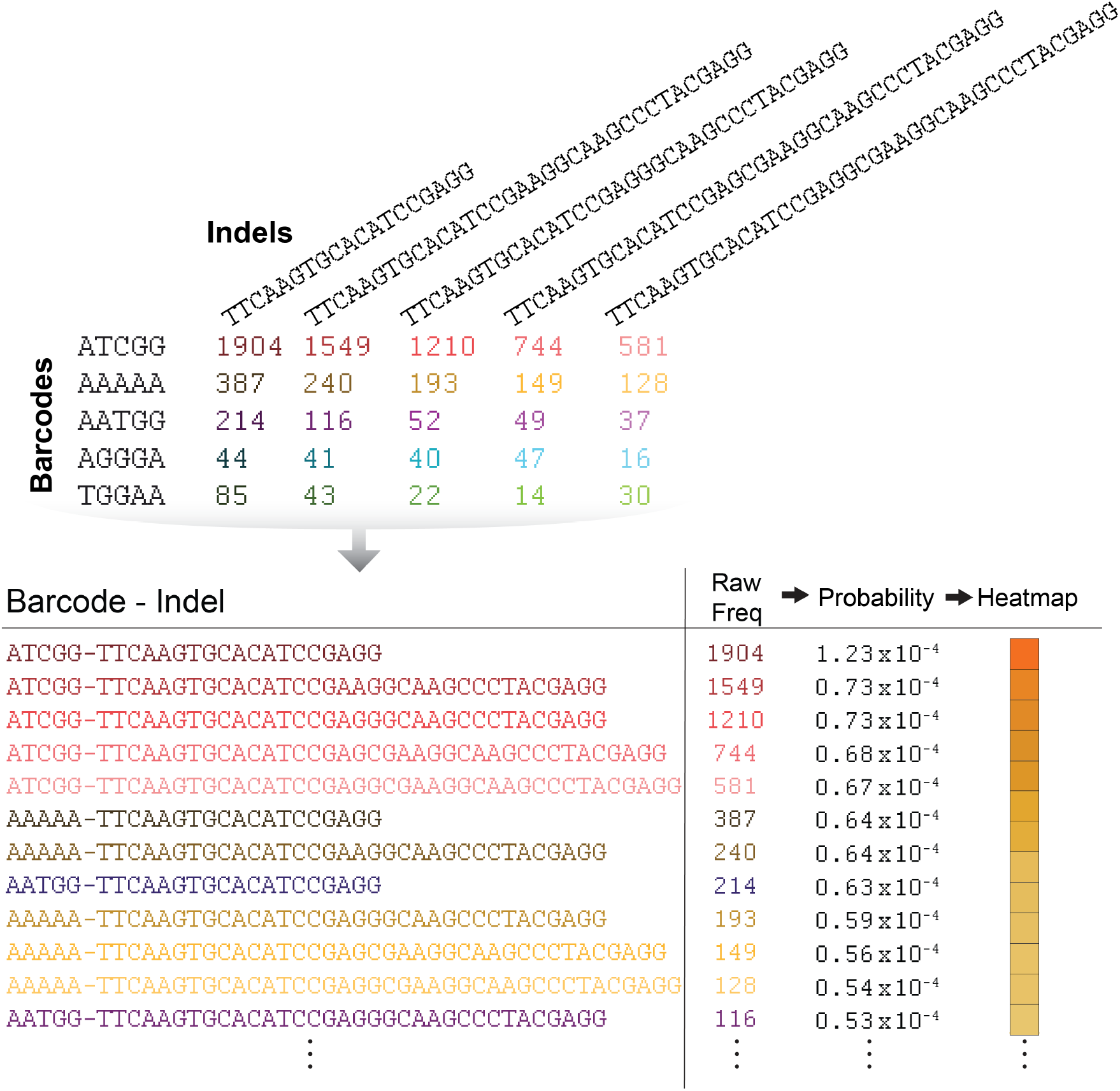
Quantitative assessment of CREAM-PUFs. To quantitatively assess the effectiveness of CREAM-PUFs, the NGS result was converted to a frequency-based array of barcode-indel combinations. The corresponding probability density functions were then calculated to enable comparison between samples.

To perform a pairwise comparison between CREAM-PUFs derived from each cell line we use a standard metric for computing distance between probability distributions, the Total Variation Distance. The results (**Figure 9**) reveal that *intra-PUF* distances (defined as the variation between a specific CREAM-PUFi.j and its corresponding repeat or freeze-thaw counterparts) are significantly smaller than *inter-PUF* distances (defined as the variation between two different CREAM-PUFs) (**Supplementary Table 12**) in all three cell lines. As an example, in HEK293 cells, for each of the two PUFi families (i={1,2}), a threshold on Total Variation Distance can be selected (i.e., 0.007 and 0.019 respectively) such that <u>all</u> *intra*-PUF distances are below-threshold (indicating a match) and <u>all</u> *inter*-PUF distances are above-threshold (indicating a no-match). Similarly, such thresholds can also be established in PUFs derived from HCT116 and HeLa cells (0.037 for HCT116 and 0.013 for HeLa, respectively). This can also be visually confirmed by contrasting *intra*-PUF color intensity (i.e., inside the red boxes of **Figure 9**) to *inter*-PUF color intensity (i.e., outside the red boxes) for each of the four PUFi families (i={1,2,3,4}).

**Fig. 9.**
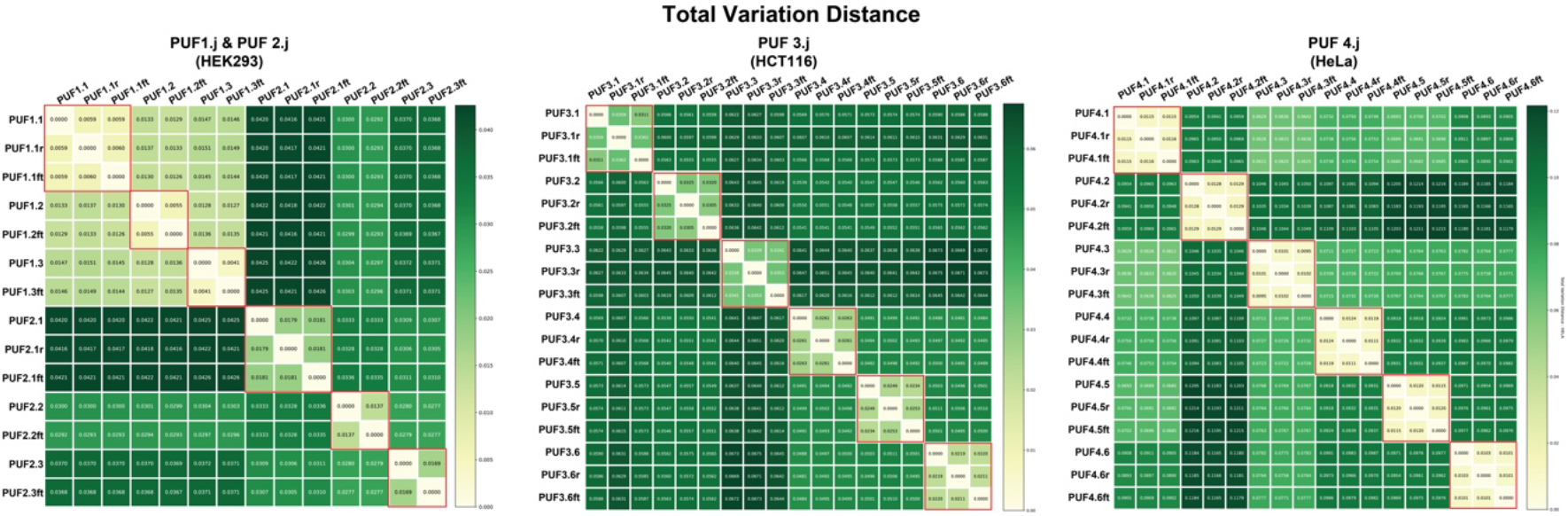
Quantitative assessment of CREAM-PUFs using total variation distance. Pairwise Total Variation Distances between all PUFs were calculated in samples derived from HEK293, HCT116, and HeLa cells.

In practice, provenance attestation can be performed quantitatively by using the Bray-Curtis dissimilarity between the end-user’s CREAM-PUF and the values stored in a database. To demonstrate the use of the Bray-Curtis in this context, we compute the *intra*-PUF and *inter*-PUF dissimilarities using the rank-ordered N most-frequent barcode-indel addresses of PUF1.1 as the reference (**Supplementary Figure 4A**). As the number of used addresses increases towards the full list (N=3478), we observe that the Bray-Curtis value between the reference (PUF1.1) and the CREAM-PUFs originating from the same Barcoded Cell Line #1 (i.e., PUF1.1r, PUF1.1ft, and PUF1.j where j={2,3}) also increases (**Supplementary Figure 4B)**. On the other hand, the Bray-Curtis value from the CREAM-PUFs originating from Barcoded Cell Line #2 (i.e., PUF2.j where j={1,2,3}) remains close to the maximum (**Supplementary Figure 4B)**. We also observe that it is possible to obtain appreciably different *intra-PUF* and *inter-PUF* values by using as few as N=10 addresses (**Supplementary Figure 4C)**. Indeed, we find it unnecessary to use the complete list, since the contribution of additional barcode-indel addresses to the difference between *intra*-PUF and *inter*-PUF Bray-Curtis dissimilarities diminishes as N increases. Overall, we observe that Bray-Curtis dissimilarity calculation using approximately 15% of the barcode-indel addresses (for all cell lines) results in lists that can provide an indisputable identification signature (**Supplementary Figures 4-9),** while being sufficiently large to prevent unauthorized reproduction, as discussed later.

Based on the above observations, we calculated the Bray-Curtis dissimilarities between all the CREAM-PUFs in each of the three cell lines, each time using PUFi,j as a reference and comparing to its repeat and freeze-thaw versions, as well as to all other CREAM-PUFs. As shown therein, a Bray-Curtis distance of 0.2 is an appropriate threshold for matching a CREAM-PUF to its repeat and freeze-thaw counterparts in HEK293-derived PUFs (**Figure 10**), while ensuring a no-match outcome when comparing to any other CREAM-PUF. For any given PUFs generated in a HEK293 cell line, the *intra-*PUF Bray-Curtis dissimilarity is never higher than 0.2, and the *inter-*PUF Bray-Curtis dissimilarities of these PUFs against those generated using the same set of barcodes (e.g., PUF1.2 vs PUF 1.3) was at least 2.6-fold higher than the corresponding *intra-*PUF Bray-Curtis dissimilarity. When compared against PUFs generated from a different set of barcodes (e.g., PUF1.2 vs PUF2.2), the difference rises to a minimum of 4.8-fold and a maximum of 12-fold increase in Bray-Curtis dissimilarity. We observe that PUFs generated using HCT116 and HeLa cells show a similar trend (**Figure 11A** and **11B**, respectively). As an example, using PUF3.1 as the reference, the *inter*-PUF Bray-Curtis dissimilarities were at least 3.4-fold higher than the corresponding *intra*-PUF dissimilarities **(Figure 11A and Supplementary Figure 10)**, a pattern which was even more pronounced in HeLa-derived PUF4.1 (> 12-fold differences between *inter-* and *intra*-PUF dissimilarities, **Figure 11B and Supplementary Figure 11**).

**Fig. 10.**
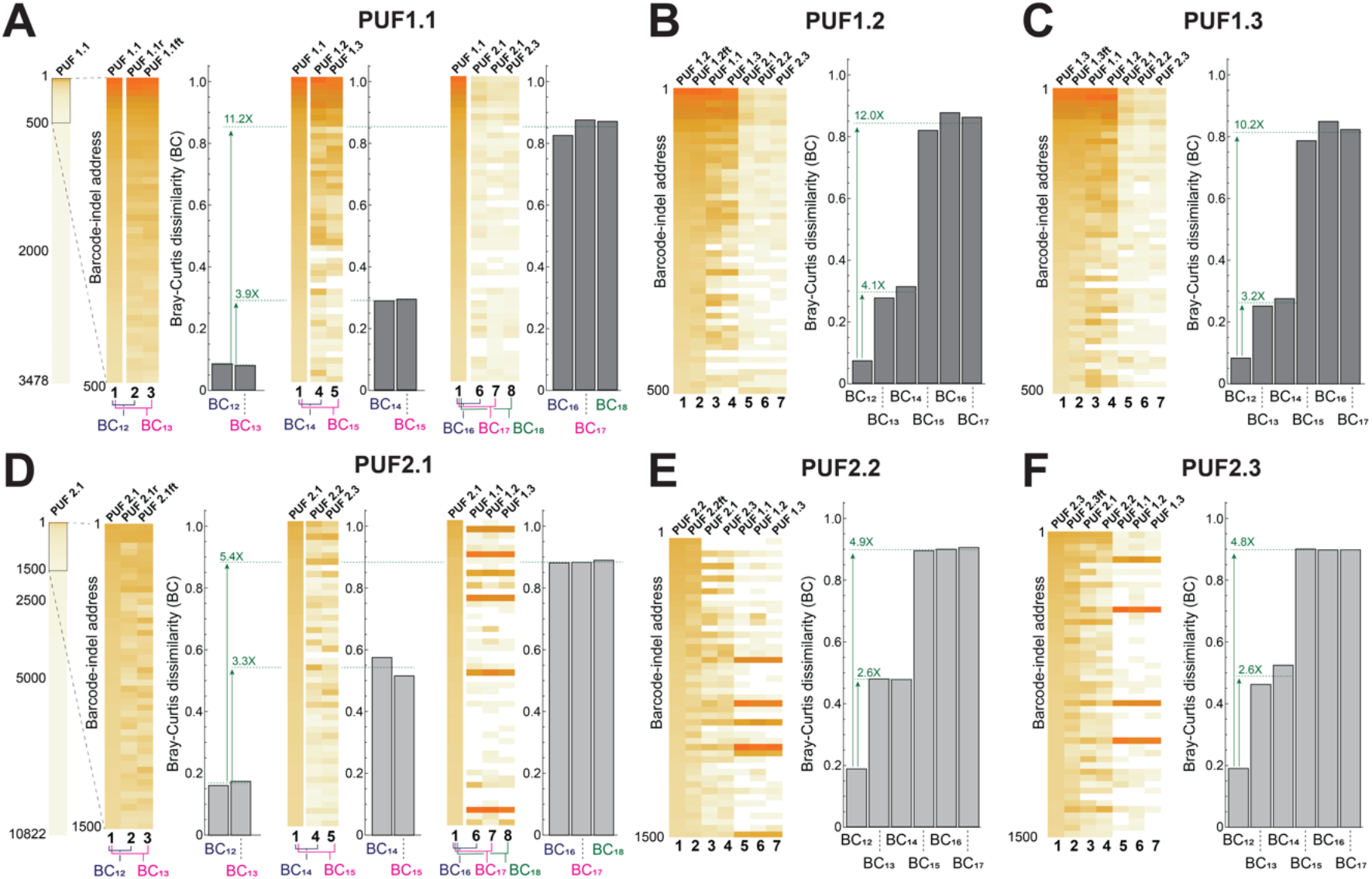
Quantitative assessment of HEK293-derived CREAM-PUFs using Bray-Curtis dissimilarity. **(A)** The difference between each pair of barcode-indel arrays is quantified using the Bray-Curtis dissimilarity method. Prior to calculating the Bray-Curtis value, the barcode-indel arrays are trimmed down to approximately 15% of the total dataset (left). Then, the Bray-Curtis dissimilarity calculations against the reference PUF (i.e., PUF1.1) are made for 3 groups: 1) technical replicates (**Left**), 2) PUFs originating from the same barcoded cell line (**Center**) and 3) PUFs originating from a different barcoded cell line (**Right**). Bray-Curtis values shown in **(B, C)** are results of an identical analysis as in **(A)** but using PUF1.2 and PUF1.3 as the reference, respectively. Bray-Curtis values shown in **(D, E and F)** are analogous results to **(A, B and C)** respectively, using PUF2.1, PUF2.2 and PUF.2.3 as the reference, respectively. Again, approximately 15% of the total dataset is used.

**Fig. 11.**
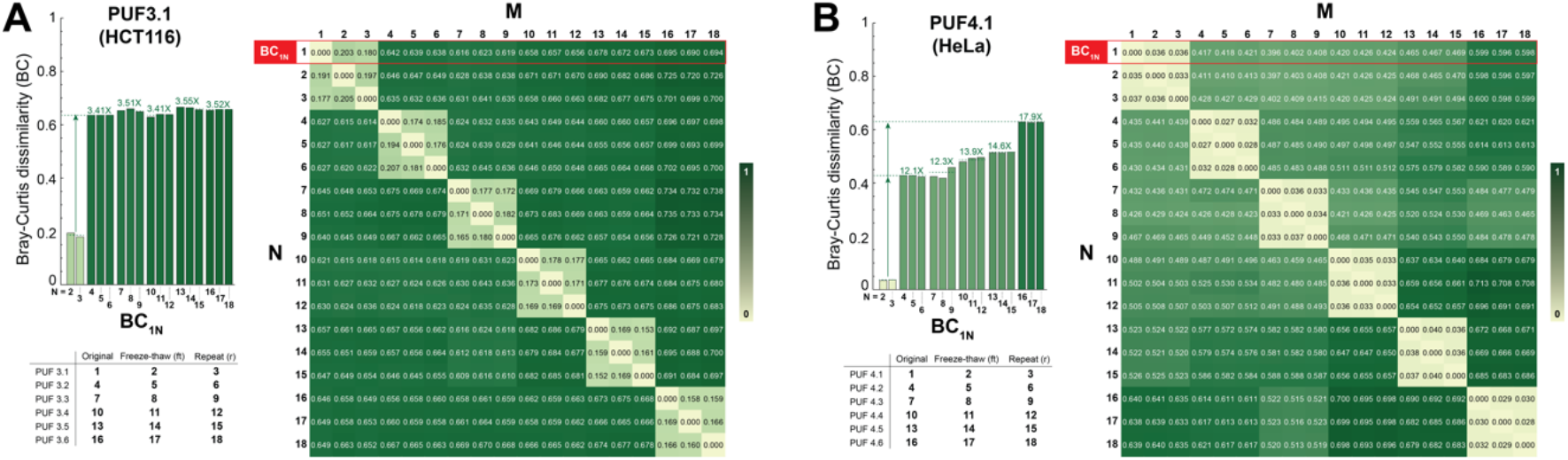
Quantitative assessment of HCT116- and HeLa-derived CREAM-PUFs using Bray-Curtis dissimilarity. **(A, Left)** Comparison of Bray-Curtis dissimilarities for a single PUF (PUF3.1) generated in HCT116 against 17 other PUFs generated in the same cell line. The barcode-indel arrays are trimmed down as described in previously in **Figure 10**. **(A, Right)** Matrix of pair-wise Bray-Curtis dissimilarity for all 18 PUFs generated in HCT116. Results shown in **(B)** are same analysis as before, with PUFs generated in HeLa cell line (PUF4.1).

We point out that a universal threshold is unnecessary, even if possible. In provenance attestation, it is sufficient to set an individual threshold for each cell line wherein a PUF has been introduced. Indeed, given a metric (e.g., Bray-Curtis dissimilarity), this threshold should be chosen to accept the signatures of all legitimately produced copies of the cell line, which the vendor stores in the CRP database, allowing a small margin to account for expected signature variation due to the freeze-thaw process or due to sequencing error, as further explained below. By individually setting this threshold for each cell line, we can optimize its ability to differentiate between PUF signatures of legitimately produced copies and illegitimate clones of a cell line.

In a noise-free case, the Bray-Curtis dissimilarity would be zero for valid PUFs. In reality, this is not the case. An important consideration here is that the Bray-Curtis values depend on the quality of the sequencing data. NGS is known to have a substitution error rate of 0.1-1% per base^39^. Therefore, in addition to our repeated sequencing experiments (i.e., PUFi,jr) and to determine the worst-case Bray-Curtis dissimilarity values originating strictly from sequencing errors, for each of the reference PUFs derived from HEK293 cells we generated 100 (artificially) mutated sequences using an error rate of 1% per base. Subsequently, the Bray-Curtis values between these mutated sequences and their PUF references were calculated using the rank-ordered barcode-indel addresses of the reference. Using these simulations, we calculated the upper bound for the Bray-Curtis dissimilarity for “valid” PUFs (**Supplementary Figure 12**). As shown in **Supplementary Table 13**, the simulated worst-case dissimilarity values accurately match a CREAM-PUF to its repeat and freeze-thaw counterparts, while ensuring a no-match outcome when comparing to any other CREAM-PUF. We note that the simulated worst-case dissimilarity values are different among PUF samples. This is expected because the underlying barcode distributions prior to applying the CRISPR-induced NHEJ are different and the absolute Bray-Curtis value depends on the average length of the sequencing reads (**Supplementary Section, ‘Bray-Curtis and sequencing reads’**).

As described earlier in **Figure 2B**, barcodes alone do not satisfy the properties required to qualify as a PUF. To validate this claim, we stably integrated a 5-nucleotide barcode library into the *AAVS1* locus of HEK293 cells in 6 parallel trials (BARCODE1-6, **Supplementary Table 14**), and subjected the samples to the two independent NGS-based amplicon sequencings. The overall barcode distribution patterns were strikingly similar among the repeats (**Supplementary Figure 13A**). Next, the Bray-Curtis dissimilarities between a BARCODEi and its sequencing repeat (BARCODEir), as well as between two distinct samples, were calculated as before **(Supplementary Figure 13B)**. As shown in **Supplementary Table 15**, the *intra*-PUF dissimilarities generally overlapped with those of *inter*-PUFs (as an example, the Bray-Curtis dissimilarity between BARCODE2 and BARCODE2r was 0.013, which was higher than the Bray-Curtis dissimilarity between BARCODE3 and BARCODE4, which was 0.011). These results confirmed our conjecture that barcodes alone do not satisfy the uniqueness requirement and therefore are not suitable to be used as PUFs.

To further investigate the uniqueness of our generated PUFs we performed additional computational analysis. Specifically, we tested whether the observed distribution of the barcode-indel addresses represents a unique combination of barcodes and indels that cannot be replicated. To achieve this, we randomly sampled a barcode sequence and an indel sequence from each of the reference HEK293-derived PUFs’ probability distribution functions, and subsequently concatenated these two sequences to generate novel combinations of barcode-indel addresses (**Supplementary Figure 14**). The same number of concatenated addresses as in the original PUF was simulated to form a novel “resampled” PUF. Specifically, for each reference PUF, 100 resampled PUFs were generated. Next, the Bray-Curtis values between these simulated sequences and their PUF references were calculated (**Supplementary Table 16**). As shown in **Supplementary Figure 15**, for all reference CREAM-PUFs, the simulated *inter-*PUF dissimilarities (i.e., Bray-Curtis values between a reference and its reshuffled samples) are between 2.8x and 3.7x larger than *intra*-PUF dissimilarities (i.e., Bray-Curtis values between a reference and its repeat or freeze-thaw counterparts), and additionally, are all larger than the worst-case dissimilarity values identified in our earlier analysis.

Collectively, these additional computational and experimental results confirm that CREAM-PUFs satisfy both the robustness and the uniqueness criteria required for serving as a cell-line provenance attestation mechanism. We further posit that CREAM-PUFs are also virtually impossible to replicate, thus unclonable. In the electronics industry, uniqueness and unclonability go hand-in-hand because silicon PUFs are inherent byproducts of the randomness of semiconductor manufacturing. Even if the PUF function is known, manufacturing an exact clone is impossible. In biology, counterfeiting a CREAM-PUF whose barcode-indel matrix is known would require DNA synthesis and integration of each individual sequence into a target cell line, followed by mixing the monoclonal cell populations to achieve the desired CREAM-PUF frequencies. While gene synthesis is becoming cheaper and synthesizing each individual fragment is feasible, integration, single cell isolation, mixing at desired proportions and, finally, validation requires prohibitive resource and time investment (**Supplementary Section, ‘Reverse Engineering a CREAM-PUF’**). Notably, the key determinants of synthesis costs and complexity (i.e., distance between the barcode and indel location and the number of barcode/indel combinations respectively) are dictated by the CREAM-PUF owner.

## Discussion

To summarize, herein we introduce a novel methodology that can be used to establish a provenance attestation protocol for commercial distribution of cell lines. Specifically, we exploit both the complexity of barcode libraries and the inherent stochasticity of DNA error-repair induced via genome editing to introduce physical unclonable functions in human cells. As valuable cell lines continue to emerge, provenance attestation to protect the investment and intellectual property of the producing company from illegal replication and to authenticate each clients’ legitimate ownership of the purchased product is bound to become essential.

Prior to silicon PUFs, the lack of provenance attestation methods fueled a counterfeiting industry (IP theft through reverse engineering, illicit overproduction, IC recycling, remarking, etc.) resulting in an estimated^40^ annual loss of $100B by legitimate semiconductor companies. The invention of silicon PUFs has not only significantly curtailed the problem but has particularly succeeded in preventing counterfeiting of the latest cutting-edge products. Silicon PUFs were introduced for the purpose of providing a unique, robust, and unclonable digital fingerprint in each copy of a legitimately produced fabricated integrated circuit. While this digital fingerprint can be used as a key to support cryptographic algorithms, its main intent is provenance attestation of the integrated circuit.

Similarly, our methodology enables the producer of a valuable cell line to insert a unique, robust and unclonable signature in each legitimately produced copy of this cell line to support provenance attestation. Successful proliferation of such genetic PUFs can be transformative for intellectual property protection of engineered cell lines. Companies can introduce CREAM-PUFs to their cells to enable unique authorization and validation, labs across the world may use this technology as a starting point for validating point-of-source, and funding agencies and journals may require CREAM-PUFs in published documents and reports for quality control and for ensuring reproducibility.

## Methods

### Cell culture and transient transfection

The HEK293 cells (catalog number: CRL-1573), HCT116 cells (catalog number: CCL-247), and HeLa cells (catalog number: CCL-2) were acquired from the American Type Culture Collection and maintained at 37°C, 100% humidity and 5% CO2. The cells were grown in Dulbecco’s modified Eagle’s medium (DMEM, Invitrogen, catalog number: 11965–1181) supplemented with 10% Fetal Bovine Serum (FBS, Invitrogen, catalog number: 26140), 0.1 mM MEM non-essential amino acids (Invitrogen, catalog number: 11140–050), and 0.045 units/mL of Penicillin and 0.045 units/mL of Streptomycin (Penicillin-Streptomycin liquid, Invitrogen, catalog number: 15140). To pass the cells, the adherent culture was first washed with PBS (Dulbecco’s Phosphate Buffered Saline, Mediatech, catalog number: 21-030-CM), then trypsinized with Trypsin-EDTA (0.25% Trypsin with EDTAX4Na, Invitrogen, catalog number: 25200) and finally diluted in fresh medium. For transient transfection, ~300,000 cells in 1 mL of complete medium were plated into each well of 12-well culture treated plastic plates (Griener Bio-One, catalog number: 665180) and grown for 16–20 hours. All transfections were then performed using 1.75 μL of JetPRIME (Polyplus Transfection) and 75 μL of JetPRIME buffer. The transfection mixture was then applied to the cells and mixed with the medium by gentle shaking.

### Flow cytometry

48–72 hours post transfection cells from each well of the 12-well plates were trypsinized with 0.1 mL 0.25% Trypsin-EDTA at 37°C for 3 min. Trypsin-EDTA was then neutralized by adding 0.9 mL of complete medium. The cell suspension was centrifuged at 1,000 rpm for 5 min and after removal of supernatants, the cell pellets were re-suspended in 0.5 mL PBS buffer. The cells were analyzed on a BD LSRFortessa flow analyzer. CFP was measured with a 445-nm laser and a 515/20 band-pass filter, and mKate with a 561-nm laser, 610 emission filter and 610/20 band-pass filter. For data analysis, 100,000 events were collected. A FSC (forward scatter)/SSC (side scatter) gate was generated using a un-transfected negative sample and applied to all cell samples. The mKate and CFP readings from un-transfected HEK293 cells were set as baseline values and were subtracted from all other experimental samples. The normalized mKate values (mKate/CFP) were then collected and processed by FlowJo. All experiments were performed in triplicates.

### Generation of barcoded stable cells

To generate the barcoded stable cells, ~10 million of the cells were seeded onto a 10 cm petri dish. 16 hours later, the cells were transiently transfected with 1 μg of the donor plasmid (Barcode-Truncated CMV-mKate-PGK1-hygromycin resistance gene) and 9 μg of CMV-SpCas9-U6-AAVS1/sgRNA plasmid using the JetPRIME reagent (Polyplus Transfection). 48 hours later, hygromycin B (Thermo Fisher Scientific, catalog number: 10687010) was added at the final concentration of 200 μg/mL. The selection lasted ~2 weeks, after which the surviving clones were pooled to generate the polyclonal stable cells. The barcoded stable cells were further expanded and maintained in the complete growth medium containing 200 μg/mL of hygromycin.

### Next generation sequencing (NGS)-based amplicon sequencing

To determine the abundance of the barcode and indel sequences, total genomic DNA was isolated from CREAM-PUF cells transfected with CMV-SpCas9-U6-sgRNA5 using the DNeasy Blood & Tissue Kit (Qiagen, catalog number: 69504). cDNA fragments harboring both barcode and expected indel sequences were PCR amplified by using ~100 ng of the genomic DNA and primers P1 and P2, which added the 5’-overhang adapter sequence P12 and the 3’-overhang adapter sequence P13 for subsequent Illumina NGS amplicon sequencing. The PCR conditions were: first one cycle of 30 s at 98°C, followed by 40 cycles of 10 s at 98°C, 30 s at 60°C, and 1 min at 72°C. The purified PCR products were then subjected to NGS-based amplicon sequencing (Illumina 100-bp paired end sequencing), which was performed at the Genome Sequencing Facility (GSF) at The University of Texas Health Science Center at San Antonio (UTHSCSA). 1 million individual reads were generated for each sample.

### Total Variation Distance

The total variation distance, *δ_TVD_*, between two probability measures *P* and *Q* for a countable sample space Ω is equal to the half of the *L*^1^ norm of these distributions or equivalently, half of the elementwise sum of the absolute difference of *P* and *Q,* as defined in Eq. 1.

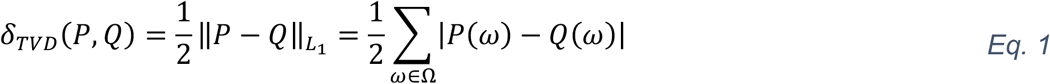

In addition, the total variation distance is the area between the two probability distribution curves defined as 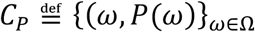 and 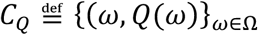. It can be shown that for a finite set Ω, the total variation distance is equal to the largest difference in probability, taken over all subsets of Ω, i.e., all possible events.

### Bray-Curtis Dissimilarity

The Bray-Curtis dissimilarity *δ_BC_* between two vectors ***u*** and ***v*** of same length ***n*** is defined in in Eq. 2:

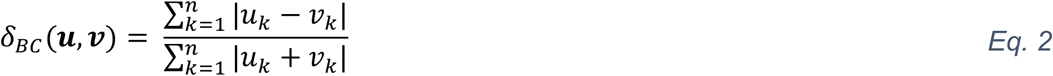

The Bray-Curtis dissimilarity has values between zero and one when all coordinates are positive.

## Acknowledgements

This work was funded by the University of Texas at Dallas SPIRE mechanism, and partially by US National Science Foundation (NSF) CAREER grant 1351354, NSF 1361355, a Cecil H. and Ida Green Endowment, and the University of Texas at Dallas.

## Author contributions

Y.L., M.B. performed research. Y.L., L.B, M.B, T.K. processed the data. All authors wrote the manuscript. L.B., Y.L., and Y.M. designed the experiments and analysis. L.B. supervised the project.

## Additional information

Supplementary information is available in the online version of the paper. Correspondence and requests for materials should be addressed to L.B.

