## Supplementary Information for "Provenance Attestation of Human Cells Using Physical Unclonable Functions"

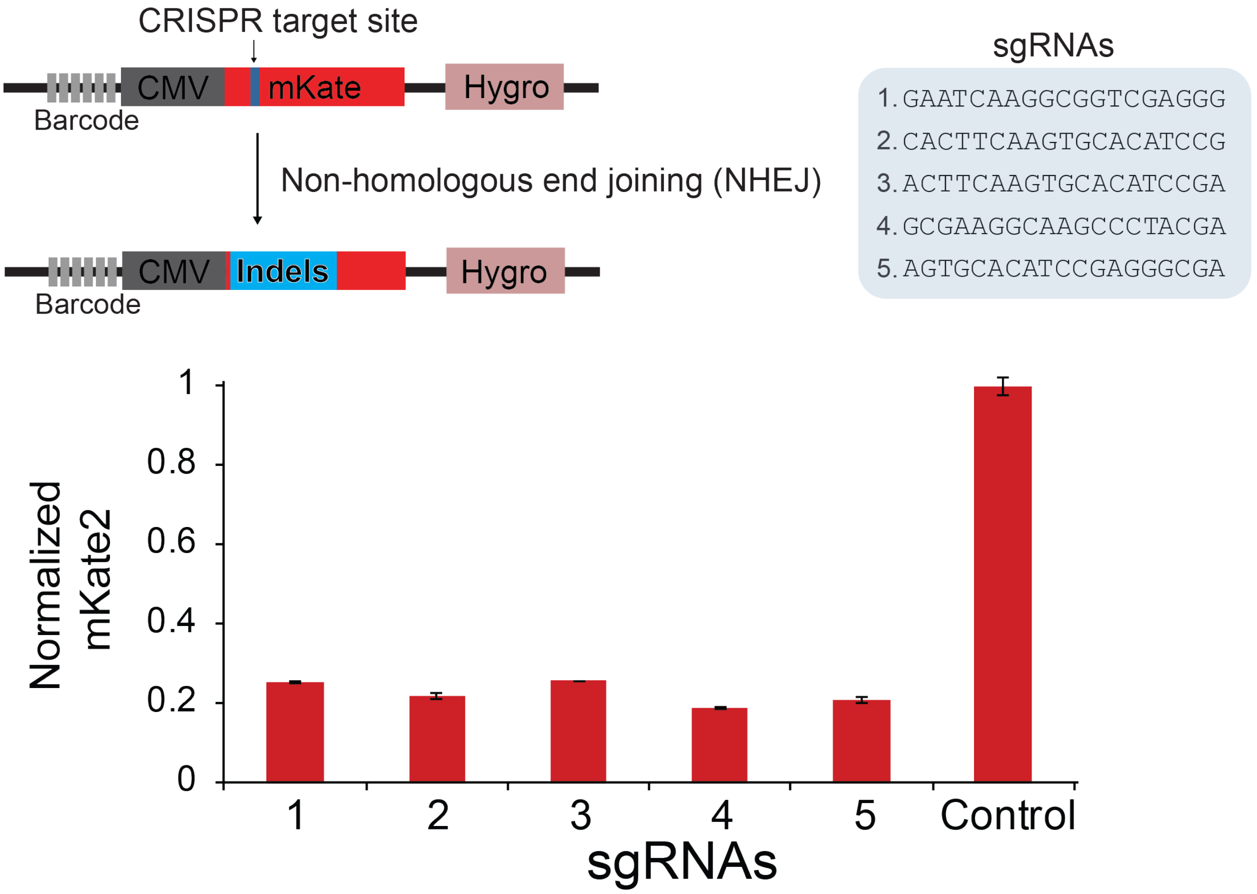


**Supplementary Figure 1. Implementation of CREAM-PUFs in HEK293 cells.** Five sgRNAs were designed to target the Open Reading Frames (ORFs) of the mKate2 construct, and demonstrated comparable efficiencies using *in vitro* fluorescence reporter assays.

A

| Sample | Barcode | Indels 1 | 2 | 3 | 4 | 5 |
|---|---|---|---|---|---|---|
| PUF3.1 | ATCGG | 102 | 97 | 156 | 82 | 84 |
| | TTCCG | 88 | 106 | 93 | 55 | 24 |
| | AGATC | 40 | 36 | 33 | 32 | 21 |
| | CGGGG | 61 | 20 | 30 | 23 | 14 |
| | ATCGC | 67 | 26 | 37 | 16 | 6 |
| PUF3.1ft | ATCGG | 118 | 74 | 56 | 36 | 37 |
| | TTCCG | 97 | 72 | 32 | 13 | 39 |
| | AGATC | 41 | 20 | 18 | 7 | 11 |
| | CGGGG | 31 | 18 | 19 | 10 | 10 |
| | ATCGC | 45 | 19 | 4 | 4 | 28 |
| PUF3.1r | ATCGG | 34 | 53 | 75 | 52 | 49 |
| | TTCCG | 56 | 75 | 57 | 29 | 10 |
| | AGATC | 37 | 19 | 23 | 15 | 10 |
| | GGGTC | 24 | 10 | 14 | 10 | 6 |
| | CGGGG | 25 | 20 | 18 | 18 | 9 |

Indel 1: CCTCGTAGGGCTTGCCTTCGCTCGGATGTGCACTTGAA
Indel 2: CCTCGTAGGGCTTGCCTTCCCCTCGGATGTGCACTTGAA
Indel 3: CCTCGTAGGGCTTGCCTTCGGATGTGCACTTGAA
Indel 4: CCTCGTAGGGCTTGCCCTCGGATGTGCACTTGAA
Indel 5: CCTCGTAGGGCTTGCCTTCGCCTCGGATGTGCACTTGAA

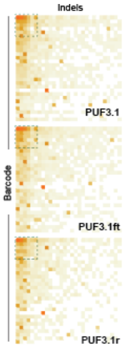


B

| Sample | Barcode | Indels 1 | 2 | 3 | 4 | 5 |
|---|---|---|---|---|---|---|
| PUF3.2 | ATCGG | 115 | 79 | 95 | 59 | 33 |
| | TTCCG | 126 | 69 | 74 | 34 | 36 |
| | CTTGG | 38 | 25 | 25 | 17 | 21 |
| | GTCGC | 29 | 22 | 27 | 11 | 11 |
| | ATACG | 14 | 21 | 16 | 9 | 10 |
| PUF3.2ft | ATCGG | 173 | 94 | 57 | 88 | 40 |
| | TTCCG | 119 | 77 | 47 | 44 | 73 |
| | CTTGG | 47 | 28 | 28 | 20 | 15 |
| | GTCGC | 33 | 14 | 17 | 11 | 8 |
| | GCCGT | 38 | 14 | 12 | 8 | 7 |
| PUF3.2r | ATCGG | 164 | 143 | 135 | 91 | 57 |
| | TTCCG | 212 | 122 | 94 | 62 | 57 |
| | CTTGG | 59 | 43 | 33 | 28 | 23 |
| | GTCGC | 38 | 26 | 21 | 18 | 17 |
| | GCCGT | 37 | 22 | 31 | 9 | 13 |

Indel 1: CCTCGTAGGGCTTGCCTTCGCTCGGATGTGCACTTGAA
Indel 2: CCTCGTAGGGCTTGCCTTCCCCTCGGATGTGCACTTGAA
Indel 3: CCTCGTAGGGCTTGCCTTCGGATGTGCACTTGAA
Indel 4: CCTCGTAGGGCTTGCCCTCGGATGTGCACTTGAA
Indel 5: CCTCGTAGGGCTTGCCTTCGCCTCGGATGTGCACTTGAA

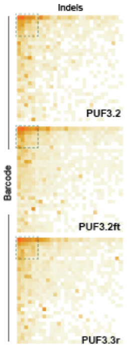


C

| Sample | Barcode | Indels 1 | 2 | 3 | 4 | 5 |
|---|---|---|---|---|---|---|
| PUF3.3 | ATCGG | 211 | 163 | 119 | 80 | 80 |
| | AATAA | 110 | 46 | 42 | 36 | 24 |
| | CACGT | 85 | 45 | 28 | 27 | 28 |
| | TAGGC | 61 | 38 | 36 | 33 | 36 |
| | TTCCG | 72 | 32 | 26 | 60 | 45 |
| PUF3.3ft | ATCGG | 204 | 150 | 103 | 78 | 89 |
| | CACGT | 69 | 49 | 35 | 25 | 29 |
| | AATAA | 93 | 43 | 34 | 28 | 28 |
| | CCGAC | 64 | 35 | 27 | 28 | 28 |
| | TTCCG | 80 | 23 | 26 | 34 | 29 |
| PUF3.3r | ATCGG | 195 | 162 | 107 | 88 | 73 |
| | CACGT | 70 | 33 | 38 | 29 | 26 |
| | AATAA | 90 | 42 | 35 | 19 | 21 |
| | TAGGC | 74 | 19 | 29 | 25 | 18 |
| | TTCCG | 62 | 27 | 26 | 36 | 20 |

Indel 1: CCTCGTAGGGCTTGCCTTCGCTCGGATGTGCACTTGAA
Indel 2: CCTCGTAGGGCTTGCCTTCCCCTCGGATGTGCACTTGAA
Indel 3: CCTCGTAGGGCTTGCCTTCGCCTCGGATGTGCACTTGAA
Indel 4: CCTCGTAGGGCTTGCCTTCGGATGTGCACTTGAA
Indel 5: CCTCGTAGGGCTTGCCCTCGGATGTGCACTTGAA

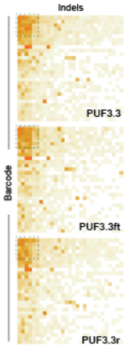


D

| Sample | Barcode | Indels 1 | 2 | 3 | 4 | 5 |
|---|---|---|---|---|---|---|
| PUF3.4 | ATCGG | 243 | 82 | 78 | 115 | 87 |
| | TTCCG | 195 | 57 | 46 | 34 | 71 |
| | CTCAA | 87 | 35 | 35 | 33 | 17 |
| | CGTAG | 67 | 33 | 24 | 30 | 18 |
| | AAACG | 56 | 26 | 23 | 12 | 13 |
| PUF3.4ft | ATCGG | 200 | 84 | 73 | 97 | 56 |
| | TTCCG | 143 | 54 | 39 | 37 | 56 |
| | CTCAA | 80 | 36 | 32 | 26 | 24 |
| | CGTAG | 65 | 19 | 14 | 24 | 17 |
| | AAACG | 41 | 16 | 18 | 11 | 13 |
| PUF3.4r | ATCGG | 192 | 71 | 64 | 92 | 86 |
| | TTCCG | 153 | 62 | 53 | 50 | 52 |
| | CTCAA | 68 | 45 | 29 | 22 | 17 |
| | AAACG | 50 | 34 | 17 | 11 | 13 |
| | CGTAG | 46 | 23 | 18 | 12 | 14 |

Indel 1: CCTCGTAGGGCTTGCCTTCGCTCGGATGTGCACTTGAA
Indel 2: CCTCGTAGGGCTTGCCTTCGGATGTGCACTTGAA
Indel 3: CCTCGTAGGGCTTGCCTTCCCCTCGGATGTGCACTTGAA
Indel 4: CCTCGTAGGGCTTGCCTTCGCCTCGGATGTGCACTTGAA
Indel 5: CCTCGTAGGGCTTGCCCTCGGATGTGCACTTGAA

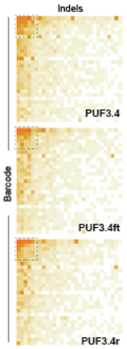


E

| Sample | Barcode | Indels 1 | 2 | 3 | 4 | 5 |
|---|---|---|---|---|---|---|
| PUF3.5 | ATCGG | 259 | 164 | 102 | 68 | 62 |
| | GGATG | 149 | 81 | 46 | 41 | 36 |
| | TGGAC | 141 | 77 | 54 | 29 | 29 |
| | TTCCG | 113 | 36 | 37 | 105 | 75 |
| | GAACA | 62 | 54 | 40 | 17 | 23 |
| PUF3.5ft | ATCGG | 227 | 153 | 94 | 57 | 41 |
| | GGATG | 138 | 66 | 51 | 32 | 53 |
| | TTCCG | 112 | 44 | 45 | 92 | 54 |
| | TGGAC | 104 | 56 | 42 | 36 | 33 |
| | ACTTG | 60 | 43 | 33 | 14 | 17 |
| PUF3.5r | ATCGG | 172 | 120 | 61 | 58 | 38 |
| | GGATG | 93 | 47 | 32 | 23 | 36 |
| | TGGAC | 73 | 30 | 42 | 25 | 22 |
| | TTCCG | 70 | 20 | 37 | 45 | 53 |
| | GAACA | 45 | 29 | 26 | 13 | 17 |

Indel 1: CCTCGTAGGGCTTGCCTTCGCTCGGATGTGCACTTGAA
Indel 2: CCTCGTAGGGCTTGCCTTCCCCTCGGATGTGCACTTGAA
Indel 3: CCTCGTAGGGCTTGCCTTCGGATGTGCACTTGAA
Indel 4: CCTCGTAGGGCTTGCCTTCGCCTCGGATGTGCACTTGAA
Indel 5: CCTCGTAGGGCTTGCCCTCGGATGTGCACTTGAA

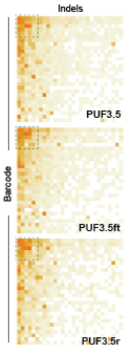


F

| Sample | Barcode | Indels 1 | 2 | 3 | 4 | 5 |
|---|---|---|---|---|---|---|
| PUF3.6 | ATCGG | 122 | 41 | 42 | 35 | 32 |
| | TTCCG | 93 | 19 | 32 | 63 | 21 |
| | TGTGG | 89 | 32 | 42 | 36 | 28 |
| | CCACT | 63 | 22 | 21 | 20 | 21 |
| | ACGTG | 45 | 16 | 9 | 22 | 34 |
| PUF3.6ft | ATCGG | 142 | 60 | 41 | 39 | 45 |
| | TTCCG | 120 | 26 | 19 | 64 | 26 |
| | TGTGG | 112 | 40 | 36 | 37 | 32 |
| | CCACT | 86 | 31 | 44 | 29 | 23 |
| | ACGTG | 44 | 24 | 21 | 15 | 51 |
| PUF3.6r | ATCGG | 193 | 63 | 48 | 51 | 45 |
| | TGTGG | 123 | 44 | 46 | 32 | 47 |
| | TTCCG | 132 | 31 | 44 | 73 | 36 |
| | CCACT | 100 | 36 | 35 | 32 | 39 |
| | TTGGG | 70 | 33 | 31 | 25 | 22 |

Indel 1: CCTCGTAGGGCTTGCCTTCGCTCGGATGTGCACTTGAA
Indel 2: CCTCGTAGGGCTTGCCTTCGGATGTGCACTTGAA
Indel 3: CCTCGTAGGGCTTGCCTTCCCCTCGGATGTGCACTTGAA
Indel 4: CCTCGTAGGGCTTGCCTTCGCCTCGGATGTGCACTTGAA
Indel 5: CCTCGTAGGGCTTGCCCTCGGATGTGCACTTGAA

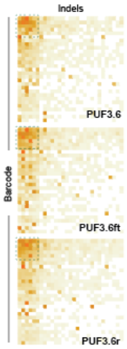


**Supplementary Figure 2. Implementation of CREAM-PUFs in HCT116 cells.** Qualitative assessment of CREAM-PUFs generated using HCT116. (A~E) Frequencies of barcode-indel addresses consisting of the 5 most commonly observed barcodes and indels (Left) and heatmap based on the same data but expanded to the top 30 most commonly observed barcodes and indels (Right) for a given PUF and its freeze-thaw counterparts and technical replicates. The green dashed square on the heatmap represents the data shown on the table. Data shown in (A) are barcode-indel addresses for PUF3.1 with their respective freeze-thaw counterpart and technical replicate. Data shown in (B~E) are for PUFs 3.2 to 3.6, respectively, which are produced identically to PUF3.1 using the same barcoded cell line and same sgRNA to introduce indels.

**A**

| Sample | Barcode | Indels 1 | 2 | 3 | 4 | 5 |
|---|---|---|---|---|---|---|
| PUF4.1 | TGGGG | 7479 | 4621 | 2735 | 2789 | 1300 |
| | CGGGG | 1033 | 135 | 88 | 96 | 635 |
| | AGGGG | 266 | 169 | 125 | 94 | 81 |
| | CGCGT | 144 | 89 | 751 | 48 | 49 |
| | ACGCC | 22 | 707 | 8 | 5 | 6 |
| PUF4.1ft | CGGGG | 7989 | 4876 | 3024 | 2907 | 1352 |
| | TGGGG | 1084 | 142 | 109 | 76 | 732 |
| | AGGGG | 246 | 202 | 138 | 98 | 103 |
| | CGCGT | 148 | 100 | 836 | 42 | 60 |
| | CACAA | 20 | 744 | 17 | 4 | 9 |
| PUF4.1r | TGGGG | 5017 | 3074 | 1936 | 1849 | 798 |
| | CGGGG | 706 | 115 | 56 | 38 | 421 |
| | CGCGT | 164 | 111 | 64 | 61 | 72 |
| | ACGCC | 102 | 65 | 534 | 31 | 33 |
| | GGGGG | 16 | 461 | 4 | 8 | 5 |

Indel 1: CCTCGTAGGGCTTGCCTTCGCCTCGGATGTGCACTTGAA
Indel 2: CCTCGTAGGGCTTGCCTTCCCCTCGGATGTGCACTTGAA
Indel 3: CCTCGGATGTGCACTTGAA
Indel 4: CCTCGTAGGGCTTGCCTTCGCTCGGATGTGCACTTGAA
Indel 5: CCTCGTAGGGCTTGCCTTCGGATGTGCACTTGAA

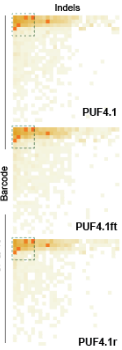


**B**

| Sample | Barcode | Indels 1 | 2 | 3 | 4 | 5 |
|---|---|---|---|---|---|---|
| PUF4.2 | CGGAG | 7681 | 5075 | 4047 | 4254 | 2991 |
| | ATAGG | 1221 | 1091 | 712 | 1023 | 120 |
| | AAGTC | 179 | 133 | 838 | 130 | 63 |
| | TCGTG | 124 | 97 | 77 | 68 | 58 |
| | AGAAG | 108 | 876 | 61 | 54 | 38 |
| PUF4.2ft | CGGAG | 7989 | 4876 | 3024 | 2907 | 1352 |
| | ATAGG | 1084 | 142 | 109 | 76 | 732 |
| | AAGTC | 246 | 202 | 138 | 98 | 103 |
| | TCGTG | 148 | 100 | 836 | 42 | 60 |
| | AGAAG | 20 | 744 | 17 | 4 | 9 |
| PUF4.2r | CGGAG | 7603 | 4992 | 4046 | 4170 | 3027 |
| | ATAGG | 1191 | 1099 | 736 | 1024 | 160 |
| | AAGTC | 168 | 149 | 792 | 100 | 75 |
| | TCGTG | 129 | 97 | 70 | 74 | 61 |
| | AGAAG | 100 | 814 | 64 | 62 | 33 |

Indel 1: CCTCGTAGGGCTTGCCTTCGCTCGGATGTGCACTTGAA
Indel 2: CCTCGTAGGGCTTGCCCTCGGATGTGCACTTGAA
Indel 3: CCTCGTAGGGCTTGCCTTCCCCTCGGATGTGCACTTGAA
Indel 4: CCTCGTAGGGCTTGCCTTCGGAA
Indel 5: CCTCGGATGTGCACTTGAA

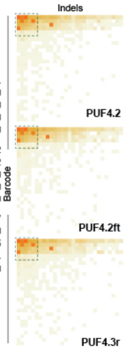


**C**

| Sample | Barcode | Indels 1 | 2 | 3 | 4 | 5 |
|---|---|---|---|---|---|---|
| PUF4.3 | CGGAG | 7298 | 4915 | 3444 | 3231 | 2379 |
| | ATAGG | 1653 | 134 | 75 | 83 | 87 |
| | AGAAG | 131 | 95 | 59 | 55 | 793 |
| | AAGTC | 271 | 149 | 109 | 88 | 94 |
| | TCGTG | 65 | 42 | 27 | 14 | 26 |
| PUF4.3ft | CGGAG | 6118 | 4276 | 2792 | 2119 | 2767 |
| | ATAGG | 1455 | 117 | 93 | 92 | 64 |
| | AGAAG | 110 | 65 | 36 | 703 | 41 |
| | AAGTC | 215 | 122 | 80 | 71 | 67 |
| | TCGTG | 54 | 40 | 19 | 27 | 19 |
| PUF4.3r | CGGAG | 4601 | 3165 | 2059 | 2018 | 1462 |
| | ATAGG | 1011 | 80 | 53 | 55 | 54 |
| | AGAAG | 73 | 42 | 23 | 32 | 473 |
| | AAGTC | 170 | 84 | 55 | 57 | 61 |
| | TCGTG | 29 | 29 | 15 | 16 | 13 |

Indel 1: CCTCGTAGGGCTTGCCTTCGCTCGGATGTGCACTTGAA
Indel 2: CCTCGTAGGGCTTGCCTTCGGATGTGCACTTGAA
Indel 3: CCTCGTAGGGCTTGCCTTCCCTCGGATGTGCACTTGAA
Indel 4: CCTCGTAGGGCTTGCCTTCGCACTTGAA
Indel 5: CCTCGTAGGGCTTGCCTTCGCGGATGTGCACTTGAA

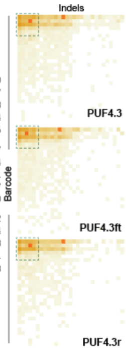


**D**

| Sample | Barcode | Indels 1 | 2 | 3 | 4 | 5 |
|---|---|---|---|---|---|---|
| PUF4.4 | CGGAG | 7419 | 1830 | 1862 | 1821 | 1250 |
| | ATAGG | 847 | 85 | 59 | 61 | 49 |
| | CGGCG | 624 | 26 | 18 | 22 | 19 |
| | AAGTC | 217 | 46 | 46 | 51 | 49 |
| | AGGAG | 23 | 6 | 7 | 4 | 5 |
| PUF4.4ft | CGGAG | 8963 | 2270 | 2154 | 2210 | 1533 |
| | ATAGG | 961 | 83 | 72 | 69 | 70 |
| | CGGCG | 843 | 23 | 41 | 24 | 24 |
| | AAGTC | 259 | 54 | 54 | 44 | 53 |
| | AGGAG | 29 | 4 | 8 | 8 | 4 |
| PUF4.4r | CGGAG | 8726 | 2149 | 2083 | 2100 | 1479 |
| | ATAGG | 869 | 74 | 81 | 66 | 66 |
| | CGGCG | 741 | 18 | 35 | 38 | 24 |
| | AAGTC | 244 | 39 | 53 | 54 | 58 |
| | AGGAG | 33 | 10 | 10 | 4 | 4 |

Indel 1: CCTCGTAGGGCTTGCCTTCGCTCGGATGTGCACTTGAA
Indel 2: CCTCGTAGGGCTTGCCTTCGGATGTGCACTTGAA
Indel 3: CCTCGTAGGGCTTGCCTTCGCGGATGTGCACTTGAA
Indel 4: CCTCGTAGGGCTTGCCTTCGGGATGTGCACTTGAA
Indel 5: CCTCGTAGGGCTTGCCTTCCCCTCGGATGTGCACTTGAA

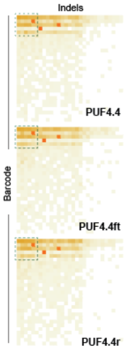


**E**

| Sample | Barcode | Indels 1 | 2 | 3 | 4 | 5 |
|---|---|---|---|---|---|---|
| PUF4.5 | CGGAG | 4958 | 4206 | 3986 | 1746 | 1492 |
| | AAGTC | 685 | 165 | 92 | 47 | 43 |
| | ATAGG | 236 | 807 | 102 | 70 | 44 |
| | AGAAG | 73 | 52 | 30 | 20 | 17 |
| | GGGTT | 589 | 11 | 8 | 8 | 3 |
| PUF4.5ft | CGGAG | 5874 | 4974 | 4678 | 2010 | 1744 |
| | AAGTC | 955 | 212 | 117 | 54 | 43 |
| | ATAGG | 272 | 1009 | 133 | 73 | 59 |
| | AGAAG | 91 | 63 | 41 | 16 | 30 |
| | GGGTT | 701 | 18 | 11 | 7 | 1 |
| PUF4.5r | CGGAG | 3099 | 2533 | 2359 | 1027 | 885 |
| | AAGTC | 453 | 110 | 79 | 31 | 22 |
| | ATAGG | 153 | 494 | 74 | 28 | 29 |
| | AGAAG | 45 | 23 | 25 | 7 | 4 |
| | GGGTT | 386 | 8 | 5 | 4 | 1 |

Indel 1: CCTCGTAGGGCTTGCCTTCGCTCGGATGTGCACTTGAA
Indel 2: CCTCGTAGGGCTTGCCTTCGGATGTGCACTTGAA
Indel 3: CCTCGTAGGGCTTGCCTTCGGTGCACTTGAA
Indel 4: CCTCGGATGTGCACTTGAA
Indel 5: CCTCGTAGGGCTTGCCTTCGTGCACTTGAA

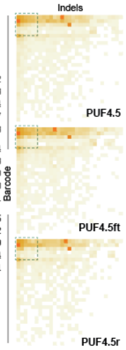


**F**

| Sample | Barcode | Indels 1 | 2 | 3 | 4 | 5 |
|---|---|---|---|---|---|---|
| PUF4.6 | CGGAG | 4217 | 4134 | 3363 | 2037 | 2696 |
| | ATAGG | 174 | 154 | 791 | 96 | 100 |
| | AAGTC | 125 | 127 | 100 | 818 | 104 |
| | AGAAG | 53 | 41 | 62 | 38 | 35 |
| | ATCGG | 33 | 31 | 21 | 16 | 11 |
| PUF4.6ft | CGGAG | 4969 | 4839 | 3844 | 2360 | 3166 |
| | ATAGG | 170 | 194 | 966 | 101 | 124 |
| | AAGTC | 142 | 149 | 141 | 925 | 88 |
| | AGAAG | 80 | 53 | 60 | 41 | 30 |
| | ATCGG | 29 | 29 | 21 | 19 | 14 |
| PUF4.6r | CGGAG | 4567 | 4592 | 3600 | 2312 | 2979 |
| | ATAGG | 190 | 175 | 866 | 121 | 104 |
| | AAGTC | 139 | 118 | 115 | 908 | 105 |
| | AGAAG | 69 | 54 | 53 | 42 | 49 |
| | ATCGG | 13 | 28 | 20 | 17 | 18 |

Indel 1: CCTCGTAGGGCTTGCCTTCGATGTGCACTTGAA
Indel 2: CCTCGTAGGGCTTGCCTTCGGATGTGCACTTGAA
Indel 3: CCTCGTAGGGCTTGCCTTCGCACTTGAA
Indel 4: CCTCGTAGGGCTTGCCCTCGGATGTGCACTTGAA
Indel 5: CCTCGCTCGGATGTGCACTTGAA

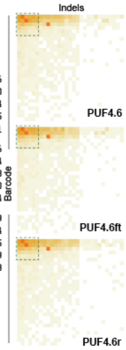


**Supplementary Figure 3. Implementation of CREAM-PUFs in HeLa cells.** Qualitative assessment of CREAM-PUFs generated using HeLa. See Supplementary Figure 2 for detailed description.

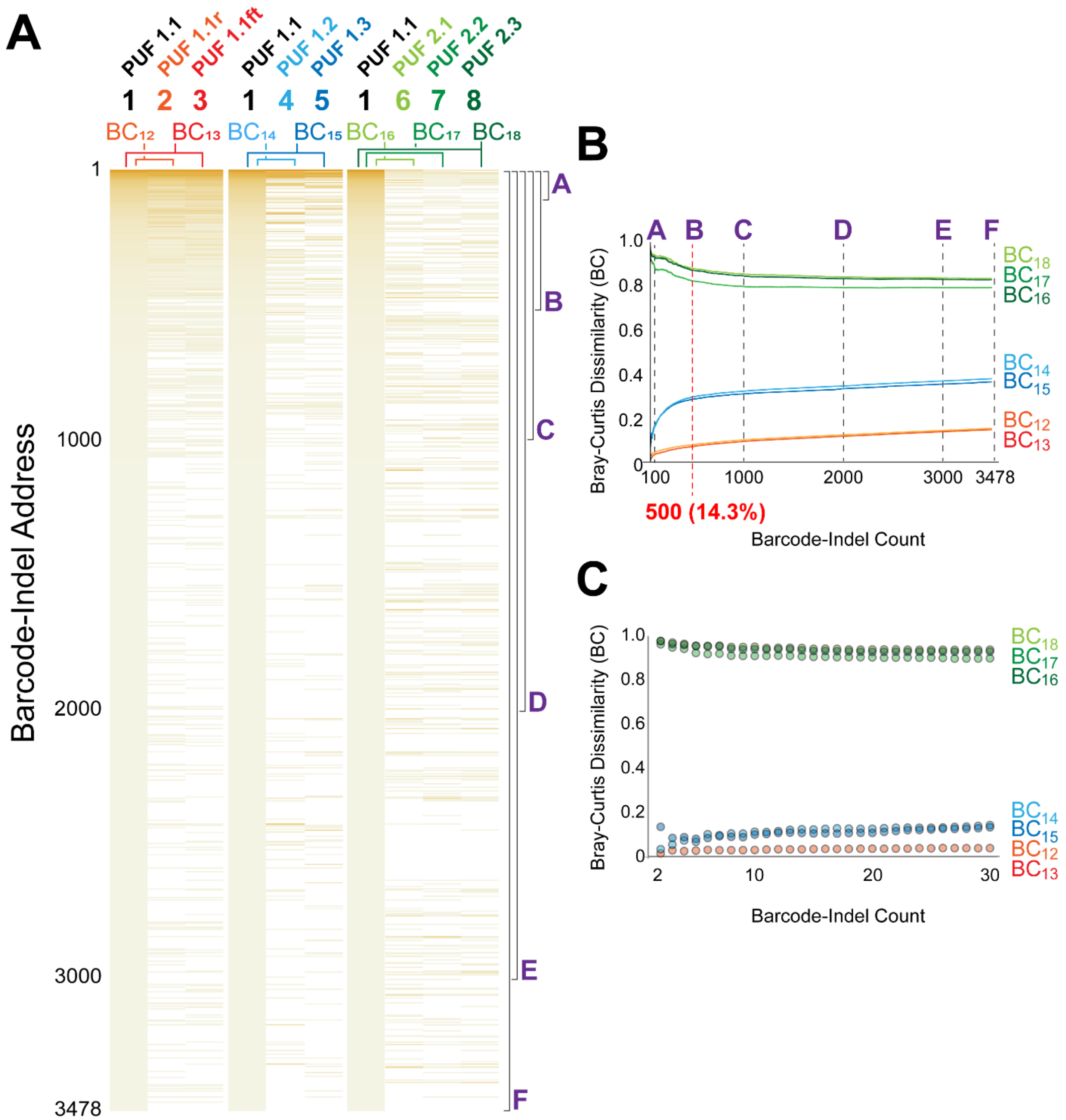


**Supplementary Figure 4. Calculation of Bray-Curtis dissimilarities using PUF 1.1 as reference with varying sampling rate. (A)** To calculate the Bray-Curtis value between 2 PUFs, the NGS results are first turned into an array of barcode-indel combinations. After sorting the array of the reference PUF based on frequency of occurrence, entries of the other arrays are then sorted to match this order. **(B)** The Bray-Curtis value between the reference and another PUF based on the size of the barcode-indel list used in the calculation, from 2 to the size of the reference sample. Purple letters indicate section of the array shown in **(A)** that corresponds to the visual representation of the list used in the calculation. The barcode-indel count shown in red indicates the list size used for analysis in the main text. **(C)** The Bray-Curtis dissimilarity based on the size of the barcode-indel list used to obtain the distance, from 2 to 30.

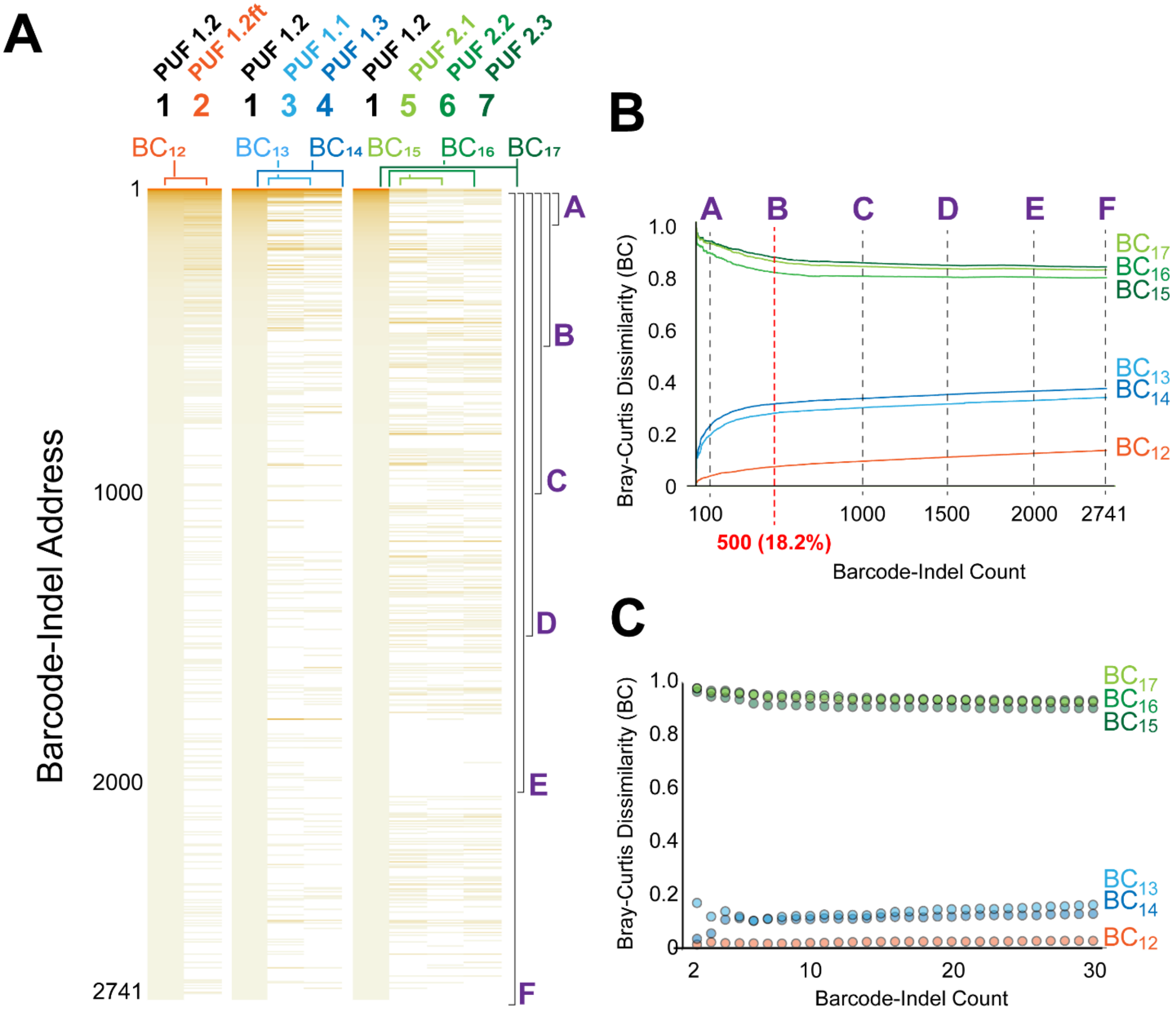


**Supplementary Figure 5. Calculation of Bray-Curtis dissimilarities using PUF 1.2 as reference with varying sampling rate.** Refer to **Supplementary Figure 4** for a detailed description.

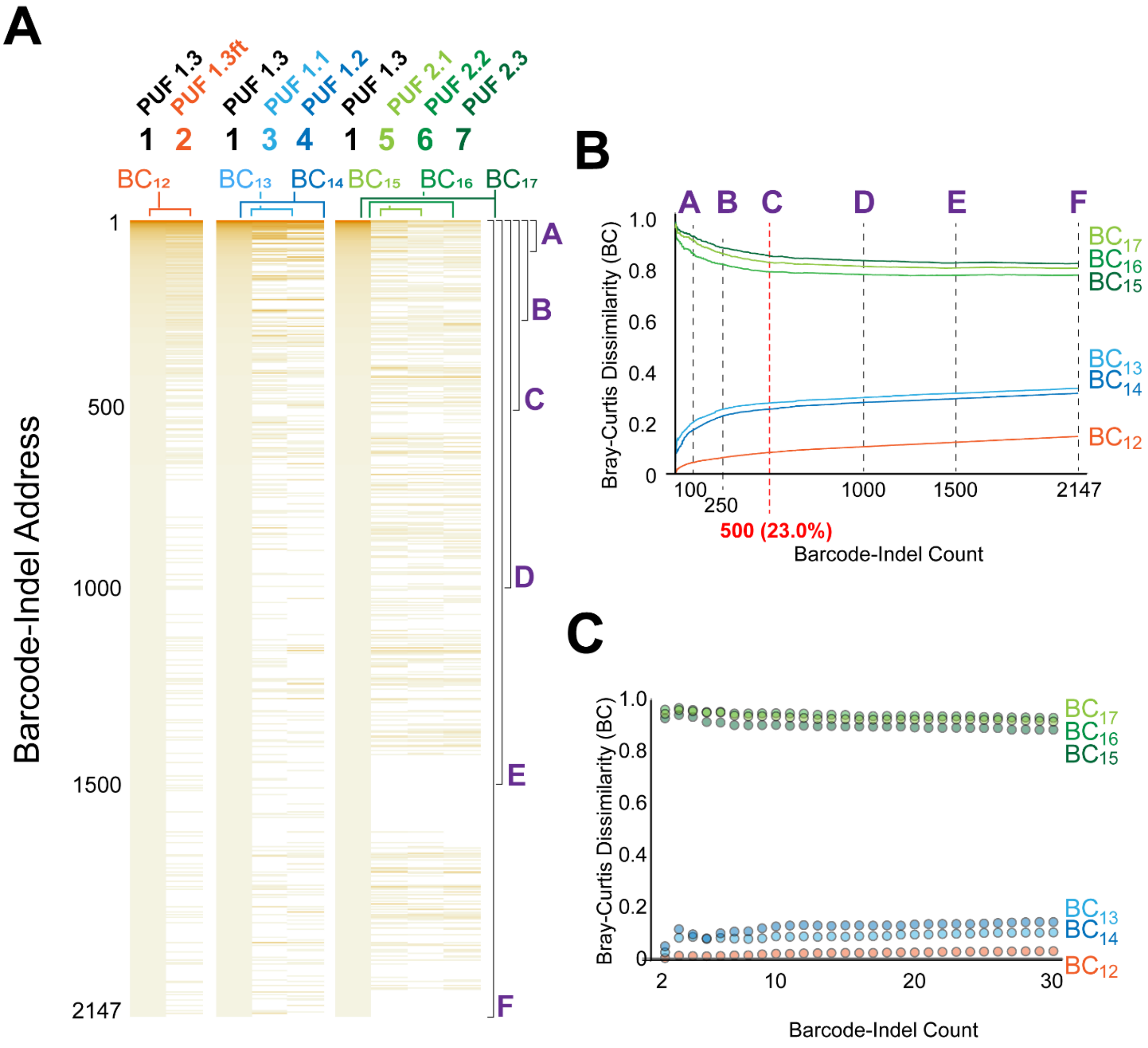


**Supplementary Figure 6. Calculation of Bray-Curtis dissimilarities using PUF 1.3 as reference with varying sampling rate.** Refer to **Supplementary Figure 4** for a detailed description.

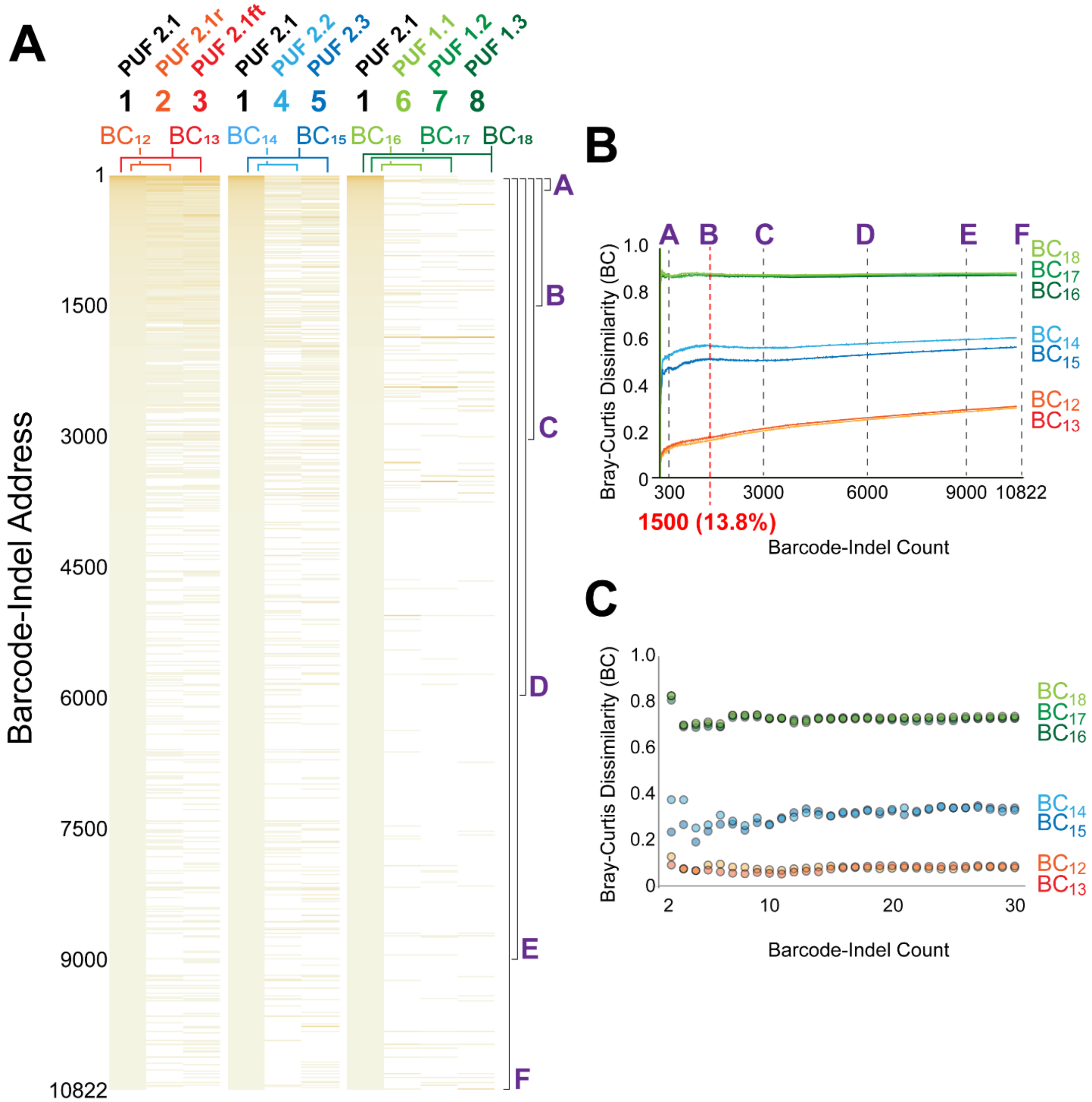


**Supplementary Figure 7. Calculation of Bray-Curtis dissimilarities using PUF 2.1 as reference with varying sampling rate.** Refer to **Supplementary Figure 4** for a detailed description.

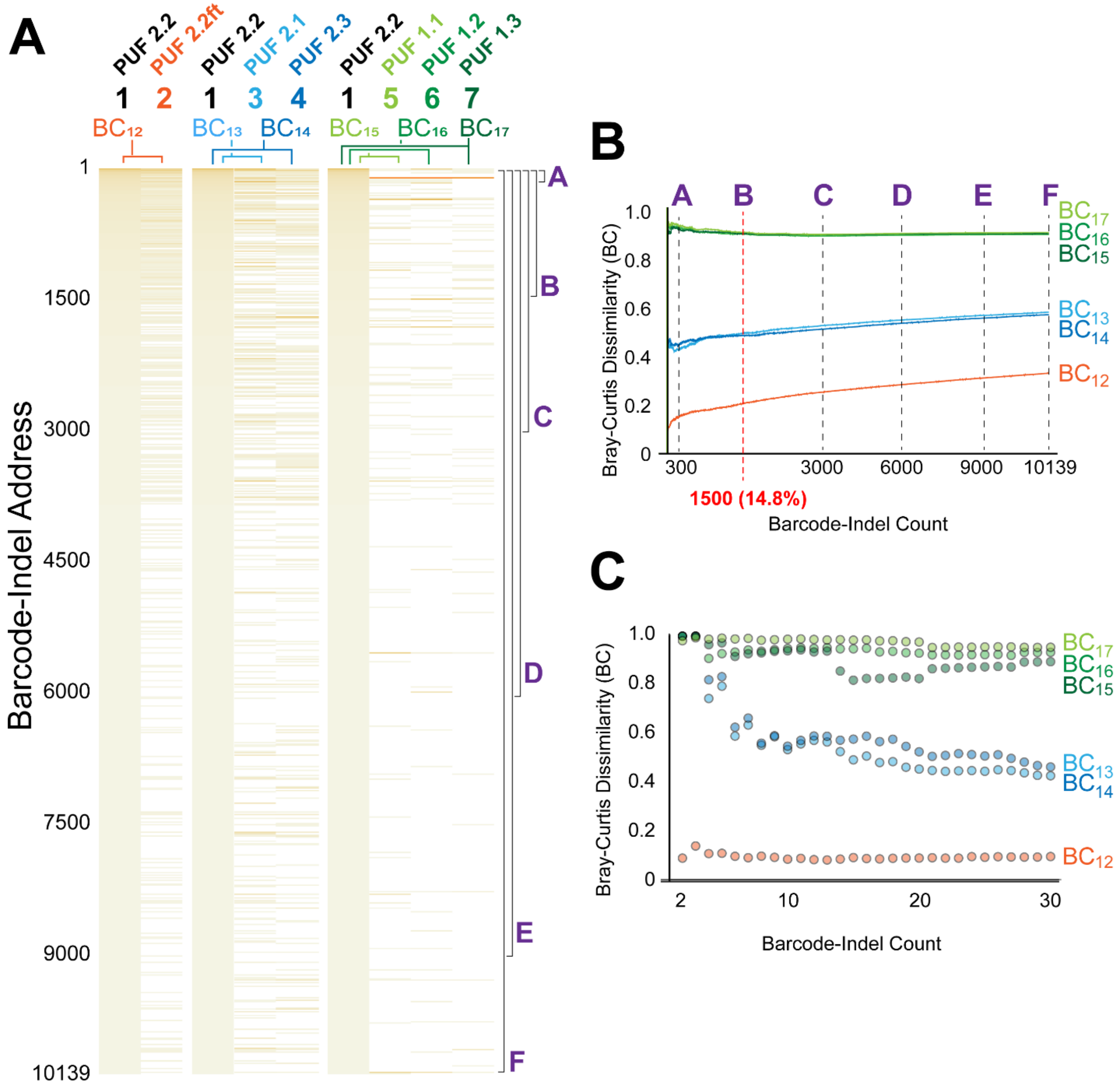


**Supplementary Figure 8. Calculation of Bray-Curtis dissimilarities using PUF 2.2 as reference with varying sampling rate.** Refer to **Supplementary Figure 4** for a detailed description.

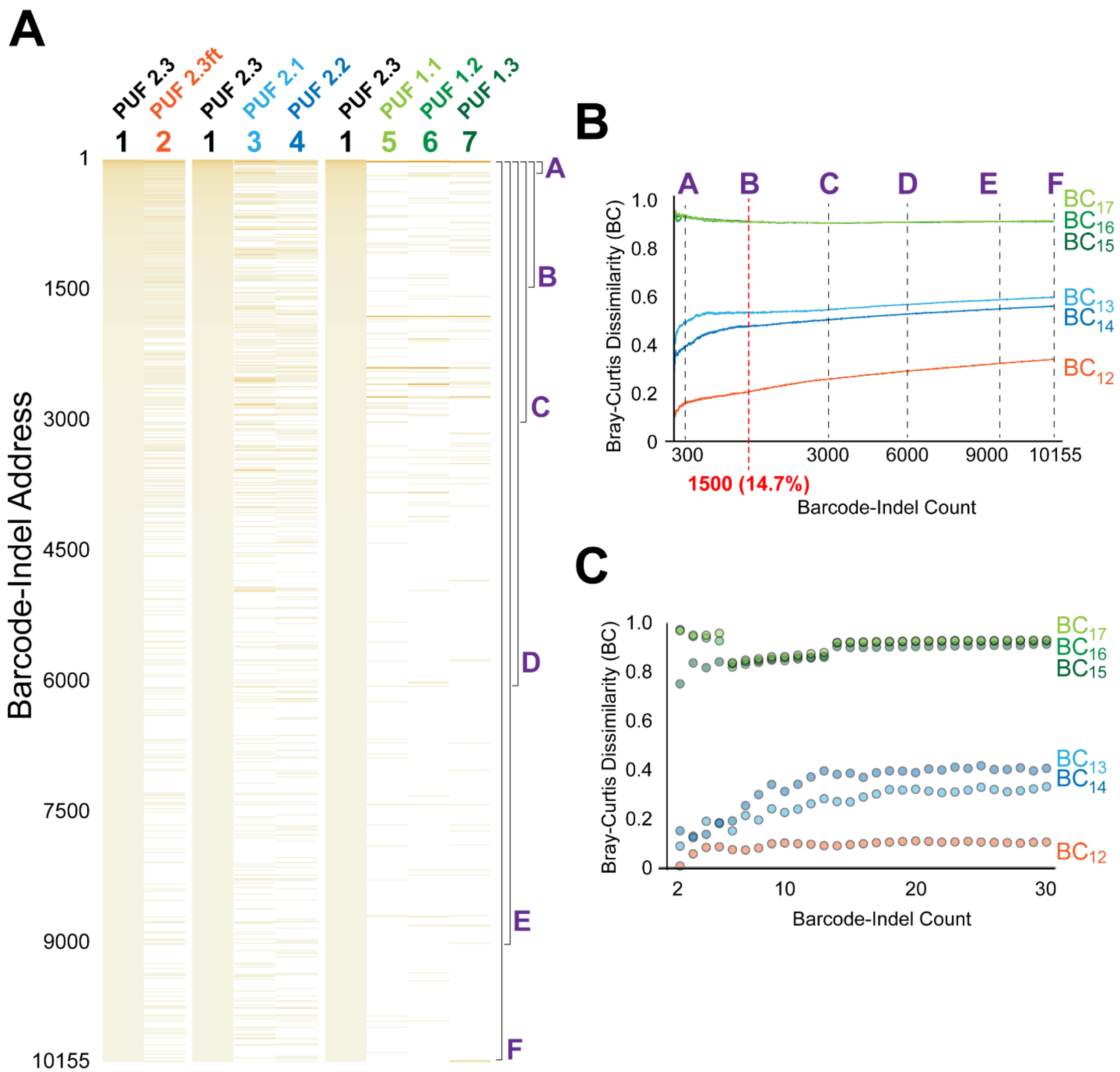


**Supplementary Figure 9. Calculation of Bray-Curtis dissimilarities using PUF 2.3 as reference with varying sampling rate.** Refer to **Supplementary Figure 4** for a detailed description.

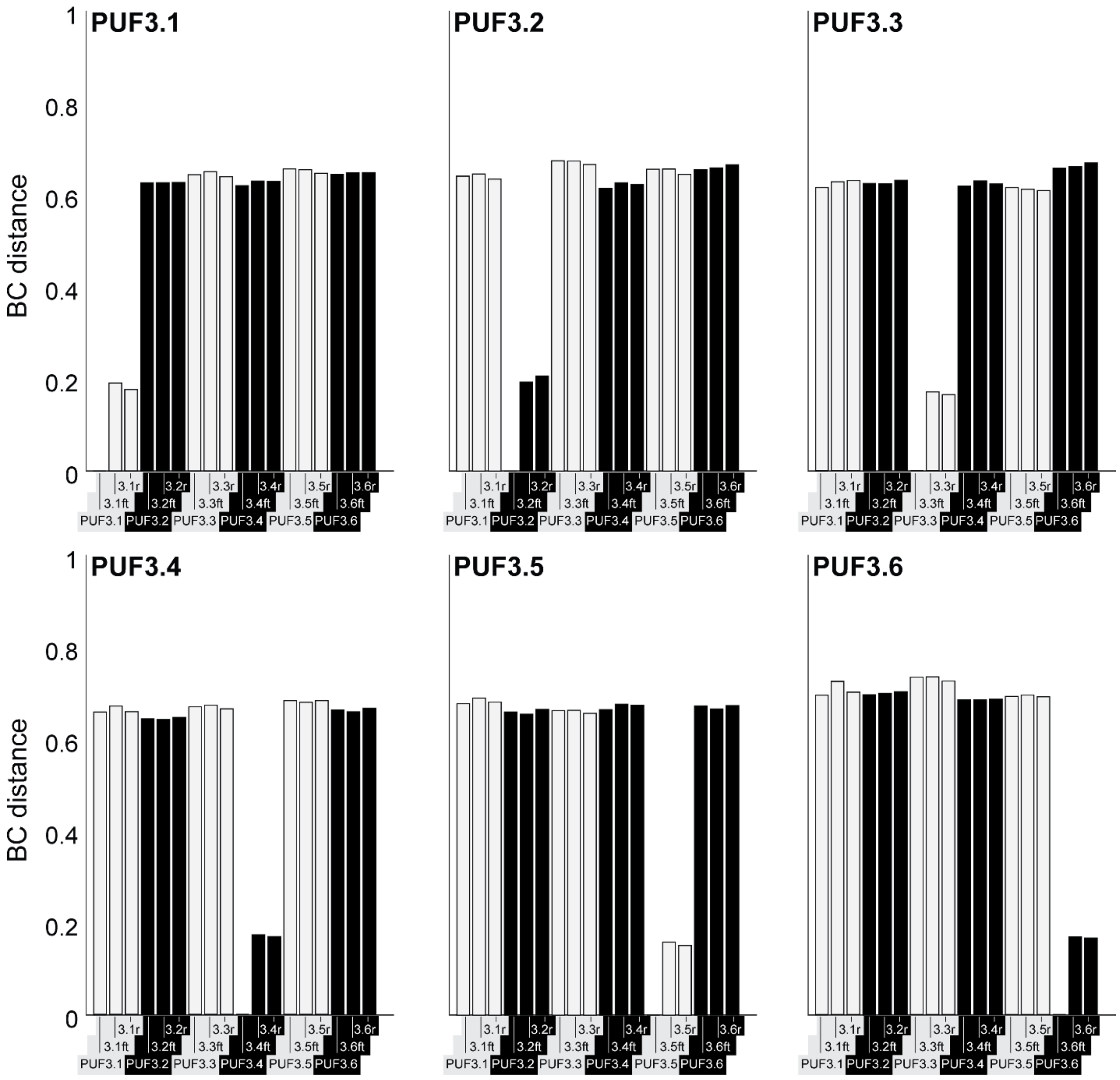


**Supplementary Figure 10. Quantitative assessment of HCT116-derived CREAM-PUFs using Bray-Curtis dissimilarity.** Comparison of Bray-Curtis dissimilarities for a single PUF3.i (i={1,2,3,4,5,6}) generated in HCT116 against 17 other PUFs generated in the same cell line.

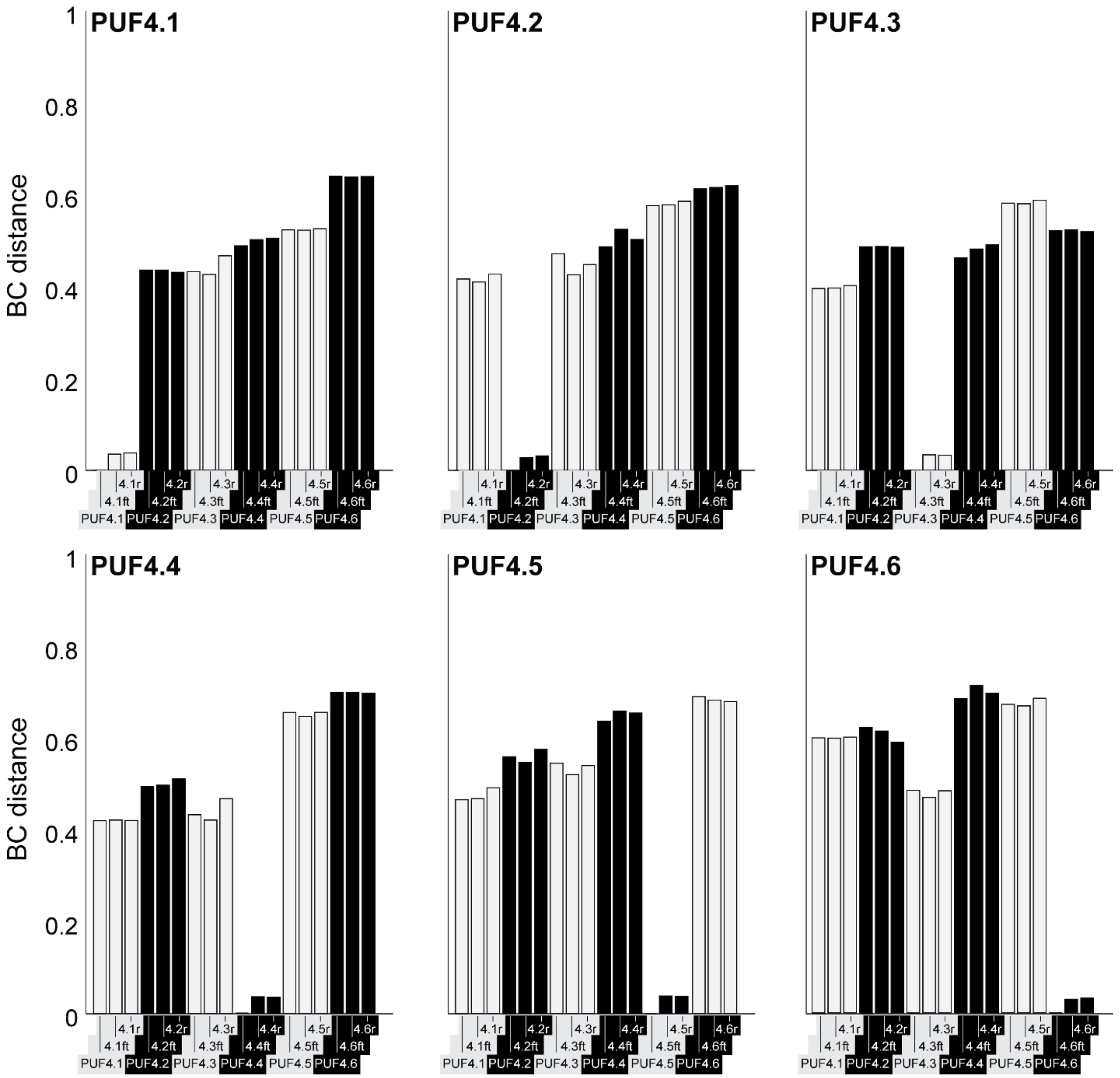


**Supplementary Figure 11. Quantitative assessment of HeLa-derived CREAM-PUFs using Bray-Curtis dissimilarity.** Comparison of Bray-Curtis dissimilarities for a single PUF4.i (i={1,2,3,4,5,6}) generated in HeLa against 17 other PUFs generated in the same cell line.

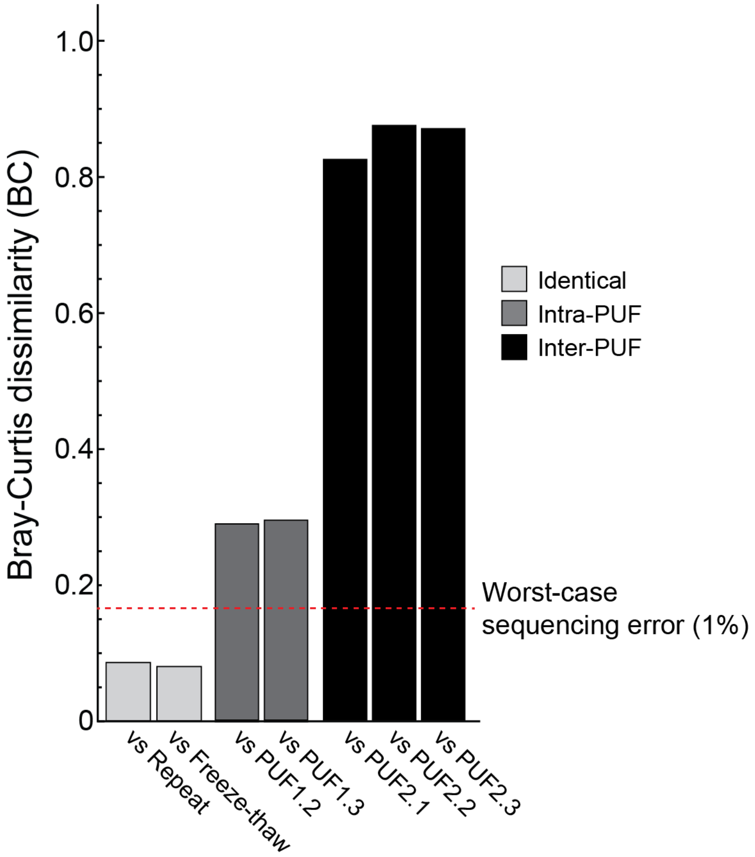


**Supplementary Figure 12. Simulated maximum Bray-Curtis dissimilarity from sequencing error for PUFs.** To obtain the worst-case Bray-Curtis values from sequencing error, each PUF barcode-indel sequencing data were mutated *in silico* using an error rate of 1% per base. The resulting dataset was then used to calculate the Bray-Curtis value against the original sequence and the technical replicates of the original sequence (repeat and freeze-thaw). The value shown for worst-case sequencing error is an average of 100 different simulations.

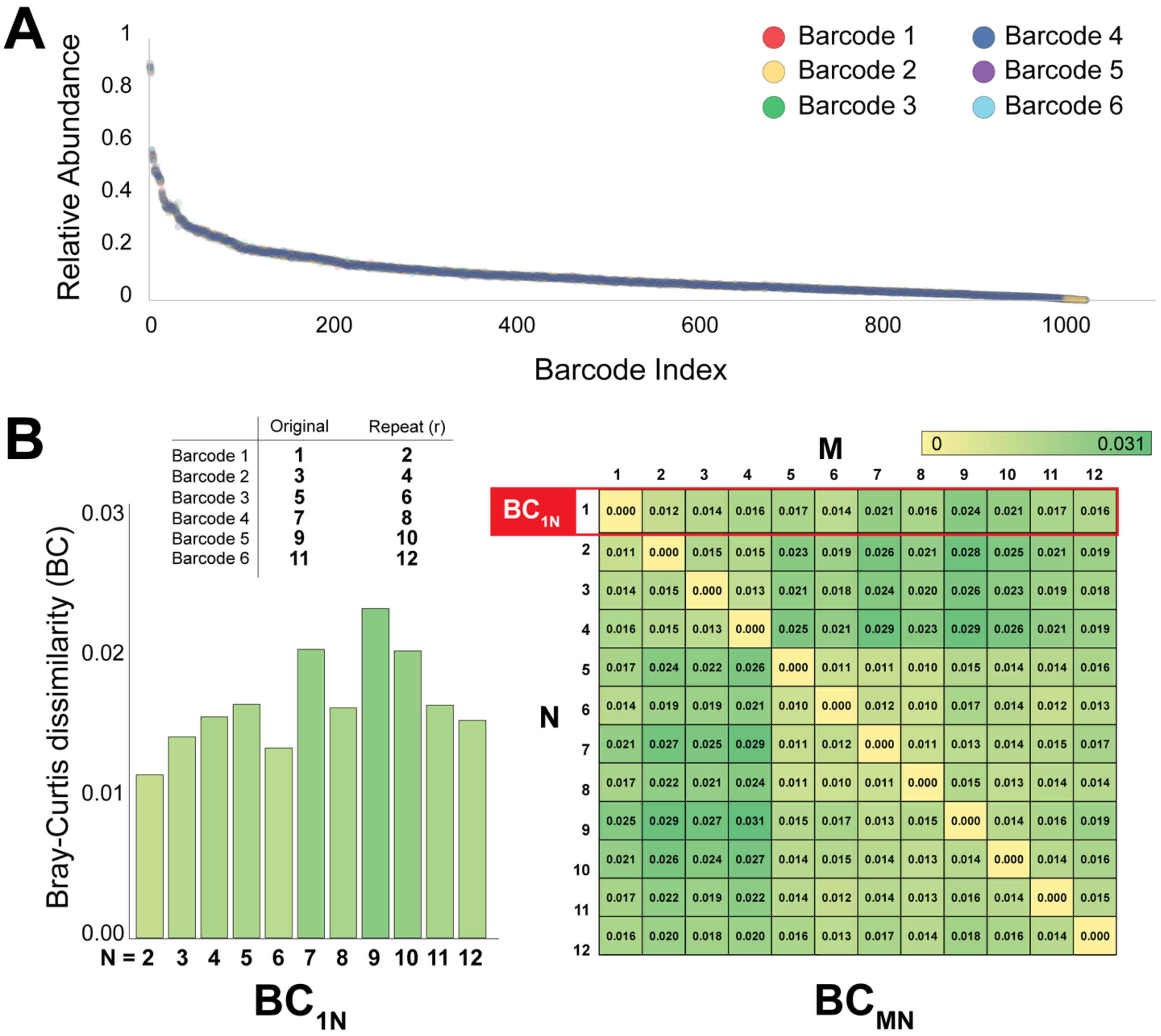


**Supplementary Figure 13. Barcode library alone does not satisfy the uniqueness requirement of PUFs.** A 5-nucleotide barcode library was stably integrated into the AAVS1 locus of HEK293 cells in 6 parallel trials. **(A)** The relative abundances of stably integrated barcodes in 6 replicates. **(B)** The Bray-Curtis dissimilarity values between barcode 1 and all other 6 samples and their NGS sequencing replicates (left) and of any given pair of all barcodes (right). Note the *intra*-sample dissimilarities generally overlapped with those of *inter*-samples, thus violating the uniqueness requirement of PUFs.

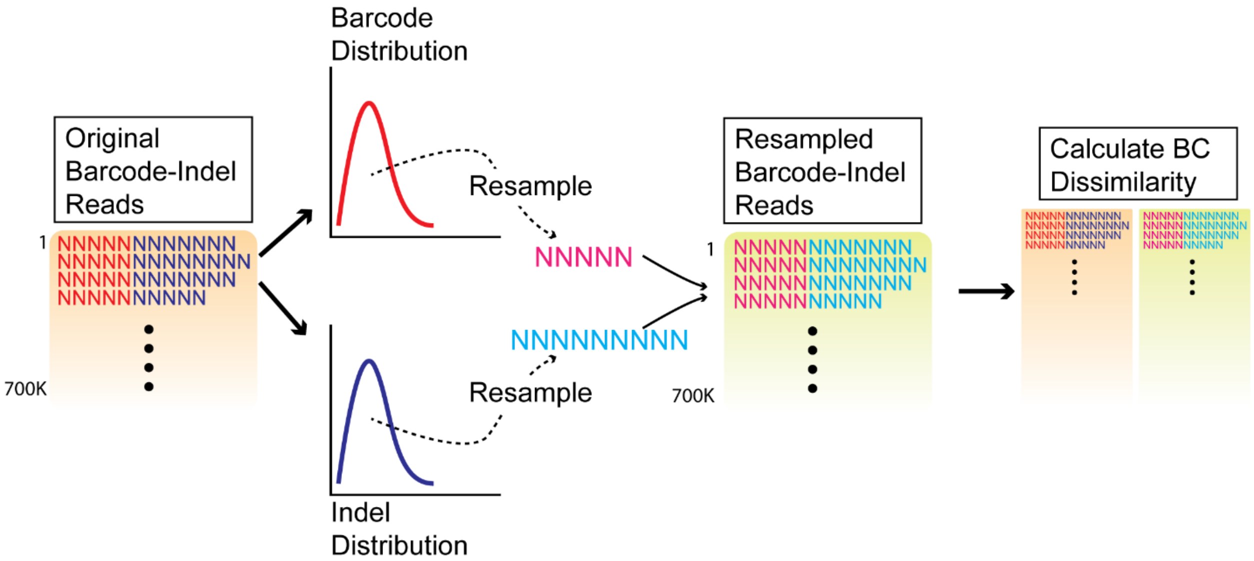


**Supplementary Figure 14. Procedure for generating resampled Barcode-Indel reads and corresponding BC dissimilarity**

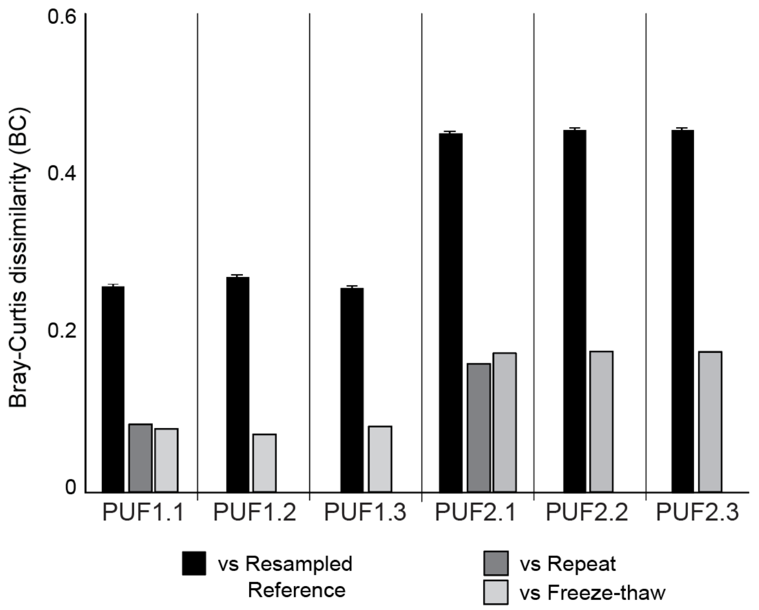


**Supplementary Figure 15. Bray-Curtis dissimilarities for intra-PUFs and simulated inter-PUFs**.

**Supplementary Material**

**General cloning protocols**

Q5 High-Fidelity 2X Master Mix (New England Biolabs) was used for all polymerase chain reactions (PCR) according to the manufacturer's protocol. All oligonucleotides were ordered from Sigma-Aldrich and were listed in **Supplementary Table 1**. The plasmids were constructed using PCR amplification, restriction digest (all restriction enzymes were ordered from New England Biolabs), and ligation with T4 DNA ligase (New England Biolabs). Gel purification and PCR purification were performed with QIAquick Gel Extraction and PCR Purification kits (Qiagen). Transformations were performed using NEB 5-alpha electrocompetent *Escherichia Coli* (New England Biolabs). The minipreps were performed using QIAprep Spin Miniprep kit (Qiagen). The final plasmids were confirmed by both restriction enzyme digestions and direct Sanger sequencings.

**DNA constructs**

**Barcode-Truncated CMV-mKate-PGK1-hygromycin resistance gene:** CMV-mKate-PGK1-hygromycin resistance gene (unpublished results) was used as the PCR template with primers P3 and P4. The purified PCR product was then cloned into CMV-mKate-PGK1-hygromycin resistance gene vector using AscI and SbfI sites.

**CMV-SpCas9-U6-sgRNA1:** CMV-SpCas9-U6-BRIP1-sgRNA (unpublished results) was used as the PCR template with primers P5 and P6. Next, the purified PCR product was used as the PCR template with primers P5 and P7. The purified PCR product was then cloned into CMV-SpCas9 (unpublished results) vector using KpnI and XbaI sites.

**CMV-SpCas9-U6-sgRNA2:** CMV-SpCas9-U6-BRIP1-sgRNA (unpublished results) was used as the PCR template with primers P5 and P8. Next, the purified PCR product was used as the PCR template with primers P5 and P7. The purified PCR product was then cloned into CMV-SpCas9 (unpublished results) vector using KpnI and XbaI sites.

**CMV-SpCas9-U6-sgRNA3:** CMV-SpCas9-U6-BRIP1-sgRNA (unpublished results) was used as the PCR template with primers P5 and P9. Next, the purified PCR product was used as the PCR template with primers P5 and P7. The purified PCR product was then cloned into CMV-SpCas9 (unpublished results) vector using KpnI and XbaI sites.

**CMV-SpCas9-U6-sgRNA4:** CMV-SpCas9-U6-BRIP1-sgRNA (unpublished results) was used as the PCR template with primers P5 and P10. Next, the purified PCR product was used as the PCR template with primers P5 and P7. The purified PCR product was then cloned into CMV-SpCas9 (unpublished results) vector using KpnI and XbaI sites.

**CMV-SpCas9-U6-sgRNA5:** CMV-SpCas9-U6-BRIP1-sgRNA (unpublished results) was used as the PCR template with primers P5 and P11. Next, the purified PCR product was used as the PCR template with primers P5 and P7. The purified PCR product was then cloned into CMV-SpCas9 (unpublished results) vector using KpnI and XbaI sites.

**NGS (next generation sequencing)-based amplicon sequencing data analysis pipeline with sample commands**

**Step 1: extracting the 100-bp reads**

```
awk 'NR%4 ==2' < f1.fastq | cat > f2.fastq
```

```
awk 'NR%4 ==2' < r1.fastq | cat > r2.fastq
```

**Step 2: joining the paired-end reads**

```
paste -d '\0' f2.fastq r2.fastq | cat > fr1.fastq
```

**Step3: filtering out corrupted reads**

```
grep “^CTTATATTCCCAGGGCCGGTTCGCGATCGCCCTGCAGG[A-Z][A-Z][A-Z][A-Z][A-
Z]TAGTTATTAATGACTCACGGGGATTTCCAAGTCTCCACCCCATTGACGTCAATGGGACCG
CCCTCGACCGCCTTGATTCTCATGGTCTGGGTGC[A-Z]*GTGGTGGTTGTTCACGGTGCCCT”
< fr1.fastq | cat > fr2.fastq
```

**Step 4: extracting the barcode and indel sequences**

```
sed -e "s/CTTATATTCCCAGGGCCGGTTCGCGATCGCCCTGCAGG \(.*\)
TAGTTATTAATGACTCACGGGGATTTCCAAGTCTCCACCCCATTGACGTCAATGGGACCGC
CCTCGACCGCCTTGATTCTCATGGTCTGGGTGC[A-Z]*GTGGTGGTTGTTCACGGTGCCCT
[A-Z]*/\1/"  < fr2.fastq | cat > barcode1.fastq
```

```
sed -e "s/CTTATATTCCCAGGGCCGGTTCGCGATCGCCCTGCAGG[A-Z][A-Z][A-Z][A-Z][A-
Z]TAGTTATTAATGACTCACGGGGATTTCCAAGTCTCCACCCCATTGACGTCAATGGGACCG
CCCTCGACCGCCTTGATTCTCATGGTCTGGGTGC \(.*\) GTGGTGGTTGTTCACGGTGCCCT
[A-Z]* /\1/" <  fr2.fastq | cat > indel1.fastq
```

**Step 5: joining the paired barcode and indel sequences**

```
paste -d '\0' barcode1.fastq indel1.fastq | cat > fr3.fastq
```

**Step 6: isolating indels containing insertions/deletions**

```
grep -v –x '.\{45\}' fr3.fastq  | cat > fr4.fastq
```

**Reverse Engineering a CREAM-PUF**

The effort needed to reverse engineer a CREAM-PUF, i.e., to synthesize a population that produces an identical barcode-indel matrix, requires an insurmountable amount of time, effort, and cost. Indeed, doing so would necessitate that each individual barcode/indel sequence pair be individually integrated into the required cell line, followed by monoclonal verification and, ultimately, mixing of the individual cells in the right proportions to reproduce the same barcode/indel frequencies observed from the CREAM-PUF. Simply installing the barcode/indel sequence can, on average, take a single researcher up to seven attempts over 19 weeks with 472 hours of hands-on time and approximately $18,000 to complete a single CRISPR editing workflow[1], i.e., generation of the desired monoclonal cell line. Furthermore, outsourcing a CRISPR-mediated genetic knock-in, such as a barcode/indel sequence described in our CREAM-PUFs, can have a starting price of $18,000-$25,000[2,3] with a similar time of completion. This process would simply produce cells with the same barcode/indel sequences contained in an individual CREAM-PUF. For example, to replicate PUF1.1, one would need to create 500 cell lines, which would cost at least $9 million. Moreover, to dial in the right frequency of engineered cells to reproduce the CREAM-PUF, would largely be trial and error with no guarantee that it is even possible.

**Barcode-Truncated CMV-mKate-PGK1-hygromycin resistance gene Sequence**

TAGGGGTTCCGCGCACATTTCCCCGAAAAGTGCCACCTGGCCAGCTCCCATAGCTCAGTC
TGGTCTATCTGCCTGGCCCTGGCCATTGTCACTTTGCGCTGCCCTCCTCTCGCCCCCGAG
TGCCCTTGCTGTGCCGCCGGAACTCTGCCCTCTAACGCTGCCGTCTCTCTCCTGAGTCCG
GACCACTTTGAGCTCTACTGGCTTCTGCGCCGCCTCTGGCCCACTGTTTCCCCTTCCCAG
GCAGGTCCTGCTTTCTCTGACCTGCATTCTCTCCCCTGGGCCTGTGCCGCTTTCTGTCTGC
AGCTTGTGGCCTGGGTCACCTCTACGGCTGGCCCAGATCCTTCCCTGCCGCCTCCTTCAG
GTTCCGTCTTCCTCCACTCCCTCTTCCCCTTGCTCTCTGCTGTGTTGCTGCCCAAGGATGC
TCTTTCCGGAGCACTTCCTTCTCGGCGCTGCACCACGTGATGTCCTCTGAGCGGATCCTC
CCCGTGTCTGGGTCCTCTCCGGGCATCTCTCCTCCCTCACCCAACCCCATGCCGTCTTCA
CTCGCTGGGTTCCCTTTTCCTTCTCCTTCTGGGGCCTGTGCCATCTCTCGTTTCTTAGGAT
GGCCTTCTCCGACGGATGTCTCCCTTGCGTCCCGCCTCCCCTTCTTGTAGGCCTGCATCAT
CACCGTTTTTCTGGACAACCCCAAAGTACCCCGTCTCCCTGGCTTTAGCCACCTCTCCATC
CTCTTGCTTTCTTTGCCTGGACACCCCGTTCTCCTGTGGATTCGGGTCACCTCTCACTCCT
TTCATTTGGGCAGCTCCCCTACCCCCCTTACCTCTCTAGTCTGTGCTAGCTCTTCCAGCCC
CCTGTCATGGCATCTTCCAGGGGTCCGAGAGCTCAGCTAGTCTTCTTCCTCCAACCCGGG
CCCCTATGTCCACTTCAGGACAGCATGTTTGCTGCCTCCAGGGATCCTGTGTCCCCGAGC
TGGGACCACCTTATATTCCCAGGGCCGGTTCGCGATCGCCCTGCAGGNNNNNTAGTTATT
AATGACTCACGGGGATTTCCAAGTCTCCACCCCATTGACGTCAATGGGAGTTTGTTTTGGC
ACCAAAATCAACGGGACTTTCCAAAATGTCGTAACAACTCCGCCCCATTGACGCAAATGGG
CGGTAGGCGTGTACGGTGGGAGGTCTATATAAGCAGAGCTGGTTTAGTGAACCGACCAGC
TAAGACACTGCCACGGTCAGATCCGCTAGCGCTACCGGTCGCCACCATGGTGAGCGAGCT
GATTAAGGAGAACATGCACATGAAGCTGTACATGGAGGGCACCGTGAACAACCACCACTT
CAAGTGCACATCCGAGGGCGAAGGCAAGCCCTACGAGGGCACCCAGACCATGAGAATCA
AGGCGGTCGAGGGCGGCCCTCTCCCCTTCGCCTTCGACATCCTGGCTACCAGCTTCATGT
ACGGCAGCAAAACCTTCATCAACCACACCCAGGGCATCCCCGACTTCTTTAAGCAGTCCTT
CCCCGAGGGCTTCACATGGGAGAGAGTCACCACATACGAAGACGGGGGCGTGCTGACCG
CTACCCAGGACACCAGCCTCCAGGACGGCTGCCTCATCTACAACGTCAAGATCAGAGGGG
TGAACTTCCCATCCAACGGCCCTGTGATGCAGAAGAAAACACTCGGCTGGGAGGCCTCCA
CCGAGACCCTGTACCCCGCTGACGGCGGCCTGGAAGGCAGAGCCGACATGGCCCTGAAG
CTCGTGGGCGGGGGCCACCTGATCTGCAACTTGAAGACCACATACAGATCCAAGAAACCC
GCTAAGAACCTCAAGATGCCCGGCGTCTACTATGTGGACAGAAGACTGGAAAGAATCAAG
GAGGCCGACAAAGAGACCTACGTCGAGCAGCACGAGGTGGCTGTGGCCAGATACTGCGA
CCTCCCTAGCAAACTGGGGCACAGAGGTGGAGGAGGTTCCGGATCTCACGGCTTCCCTCC
CGAGGTGGAGGAGCAGGCCGCCGGCACCCTGCCCATGAGCTGCGCCCAGGAGAGCGGC
ATGGATAGACACCCTGCTGCTTGCGCCAGCGCCAGGATCAACGTCTCTAGATAACTGATCA
TAATCAGCCATACCACATTTGTAGAGGTTTTACTTGCTTTAAAAAACCTCCCACACCTCCCC
CTGAACCTGAAACATAAAATGAATGCAATTGTTGTTGTTAACTTGTTTATTGCAGCTTATAAT
GGTTACAAATAAAGCAATAGCATCACAAATTTCACAAATAAAGCATTTTTTTCACTGCATTCT
AGTTGTGGTTTGTCCAAACTCATCAATGTATCTTAACGCGTAAATTGGGCGCGCCCTTAAG
CTGGGACGGAGGCTTGTTTGCGAGGCCGCGGCCGGCCGAAGTTCCTATTCTCTAGAAAGT
ATAGGAACTTCTACCGGGTAGGGGAGGCGCTTTTCCCAAGGCAGTCTGGAGCATGCGCTT
TAGCAGCCCCGCTGGGCACTTGGCGCTACACAAGTGGCCTCTGGCCTCGCACACATTCCA
CATCCACCGGTAGGCGCCAACCGGCTCCGTTCTTTGGTGGCCCCTTCGCGCCACCTTCTA
CTCCTCCCCTAGTCAGGAAGTTCCCCCCCGCCCCGCAGCTCGCGTCGTGCAGGACGTGA
CAAATGGAAGTAGCACGTCTCACTAGTCTCGTGCAGATGGACAGCACCGCTGAGCAATGG

AAGCGGGTAGGCCTTTGGGGCAGCGGCCAATAGCAGCTTTGCTCCTTCGCTTTCTGGGCT
CAGAGGCTGGGAAGGGGTGGGTCCGGGGGCGGGCTCAGGGGCGGGCTCAGGGGCGGG
GCGGGCGCCCGAAGGTCCTCCGGAGGCCCGGCATTCTGCACGCTTCAAAAGCGCACGTC
TGCCGCGCTGTTCTCCTCTTCCTCATCTCCGGGCCTTTCGACCTGCATCCATCTAGATCTC
GATCGAGCAGCTGAAGCTTACCGCAGGCTATGAAAAAGCCTGAACTCACCGCGACGTCTG
TCGAGAAGTTTCTGATCGAAAAGTTCGACAGCGTCTCCGACCTGATGCAGCTCTCGGAGG
GCGAAGAATCTCGTGCTTTCAGCTTCGATGTAGGAGGGCGTGGATATGTCCTGCGGGTAA
ATAGCTGCGCCGATGGTTTCTACAAAGATCGTTATGTTTATCGGCACTTTGCATCGGCCGC
GCTCCCGATTCCGGAAGTGCTTGACATTGGGGAATTCAGCGAGAGCCTGACCTATTGCAT
CTCCCGCCGTGCACAGGGTGTCACGTTGCAAGACCTGCCTGAAACCGAACTGCCCGCTGT
TCTGCAGCCGGTCGCGGAGGCCATGGATGCGATCGCTGCGGCCGATCTTAGCCAGACGA
GCGGGTTCGGCCCATTCGGACCGCAAGGAATCGGTCAATACACTACATGGCGTGATTTCA
TATGCGCGATTGCTGATCCCCATGTGTATCACTGGCAAACTGTGATGGACGACACCGTCAG
TGCGTCCGTCGCGCAGGCTCTCGATGAGCTGATGCTTTGGGCCGAGGACTGCCCCGAAG
TCCGGCACCTCGTGCACGCGGATTTCGGCTCCAACAATGTCCTGACGGACAATGGCCGCA
TAACAGCGGTCATTGACTGGAGCGAGGCGATGTTCGGGGATTCCCAATACGAGGTCGCCA
ACATCTTCTTCTGGAGGCCGTGGTTGGCTTGTATGGAGCAGCAGACGCGCTACTTCGAGC
GGAGGCATCCGGAGCTTGCAGGATCGCCGCGGCTCCGGGCGTATATGCTCCGCATTGGT
CTTGACCAACTCTATCAGAGCTTGGTTGACGGCAATTTCGATGATGCAGCTTGGGCGCAG
GGTCGATGCGACGCAATCGTCCGATCCGGAGCCGGGACTGTCGGGCGTACACAAATCGC
CCGCAGAAGCGCGGCCGTCTGGACCGATGGCTGTGTAGAAGTACTCGCCGATAGTGGAA
ACCGACGCCCCAGCACTCGTCCGAGGGCAAAGGAATAGGGGAGGCTAACTGAAGCTTCC
CGGGGGTACCAAATTCGTCGACAGATCTAACTTGTTTATTGCAGCTTATAATGGTTACAAAT
AAAGCAATAGCATCACAAATTTCACAAATAAAGCATTTTTTTCACTGCATTCTAGTTGTGGTT
TGTCCAAACTCATCAATGTATCTTATGATGTCTGCATATGGAAGTTCCTATTCTCTAGAAAGT
ATAGGAACTTCGCGGCCGCTCCCACCCGCTCGTCCCCCCGCGCACCTTTGCTAGGAGCG
GGTCGCCCATGTGGCTCTCAGGTTCTGGGTACTTTTATCTGTCCCCTCCACCCCACAGTGG
GGCCACTAGGGACAGGATTGGTGACAGAAAAGCCCCATCCTTAGGCCTCCTCCTTCCTAG
TCTCCTGATATTGGGTCTAACCCCCACCTCCTGTTAGGCAGATTCCTTATCTGGTGACACA
CCCCCATTTCCTGGAGCCATCTCTCTCCTTGCCAGAACCTCTAAGGTTTGCTTACGATGGA
GCCAGAGAGGATCCTGGGAGGGAGAGCTTGGCAGGGGGTGGGAGGGAAGGGGGGGATG
CGTGACCTGCCCGGTTCTCAGTGGCCACCCTGCGCTACCCTCTCCCAGAACCTGAGCTGC
TCTGACGCGGCCGTCTGGTGCGTTTCACTGATCCTGGTGCTGCAGCTTCCTTACACTTCCC
AAGAGGAGAAGCAGTTTGGAAAAACAAAATCAGAATAAGTTGGTCCTGAGTTCTAACTTTG
GCTCTTCACCTTTCTAGTCCCCAATTTATATTGTTCCTCCGTGCGTCAGTTTTACCTGTGAG
ATAAGGCCAGTAGCCAGCCCCGTCCTGGCAGGGCTGTGGTGAGGAGGGGGGTGTCCGTG
TGGAAAACTCCCTTTGTGAGAATGGTGCGTCCTAGGTGTTCACCAGGTCGTGGCCGCCTC
TACTCCCTTTCTCTTTCTCCATCCTTCTTTCCTTAAAGAGTCCCCAGTGCTATCTGGGACAT
ATTCCTCCGCCCAGAGCAGGGTCCCGCTTCCCTAAGGCCCTGCTCTGGGCTTCTGGGTTT
GAGTCCTTGGCAAGCCCAGGAGAGGCGCTCAGGCTTCCCTGTCCCCCTTCCTCGTCCACC
ATCTCATGCCCCTGGCTCTCCTGCCCCTTCCCTACAGGGGTTCCTGGCTCTGCTCTTCAGA
CTGAGCCCCGTTCCCCTGCATCCCCGTTCCCCTGCATCCCCCTTCCCCTGCATCCCCCAG
AGGCCCCAGGCCACCTACTTGGCCTGGACCCCACGAGAGGCCACCCCAGCCCTGTCTAC
CAGGCTGCCTTTTGGGTGGATTCTCCTCCAACTGTGGGGTGACTGCTTGGCAAACTCACC
GGTACCCGGCCGCGACTCTAGATCATAATCAGCTCGAGCCTTAACAAGCTTCGAAACGATA
TGGGCTGAATACAAAAACGATATGGGCTGAATACAAAAACGATATGGGCTGAATACAAACC
GCTTGAAGTCTTTAATTAAACCGCTTGAAGTCTTTAATTAAACCGCTTGAAGTCTTTAATTAA

AGGATCCACCGGATCTAGATAACTGATCATAATCGCGGCCGCACTCCTCAGGTGCAGGCT
GCCTATCAGAAGGTGGTGGCTGGTGTGGCCAATGCCCTGGCTCACAAATACCACTGAGAT
CTTTTTCCCTCTGCCAAAAATTATGGGGACATCATGAAGCCCCTTGAGCATCTGACTTCTG
GCTAATAAAGGAAATTTATTTTCATTGCAATAGTGTGTTGGAATTTTTTGTGTCTCTCACTCG
GAAGGACATATGGGAGGGCAAATCATTTAAAACATCAGAATGAGTATTTGGTTTAGAGTTTG
GCAACATATGCCATATGCTGGCTGCCATGAACAAAGGTGGCTATAAAGAGGTCATCAGTAT
ATGAAACAGCCCCCTGCTGTCCATTCCTTATTCCATAGAAAAGCCTTGACTTGAGGTTAGAT
TTTTTTTATATTTTGTTTTGTGTTATTTTTTTCTTTAACATCCCTAAAATTTTCCTTACATGTTT
TACTAGCCAGATTTTTCCTCCTCTCCTGACTACTCCCAGTCATAGCTGTCCCTCTTCTCTTA
TGAAGATCCCTCGACCTGCAGCCCAAGCTTGGCGTAATCATGGTCATAGCTGTTTCCTGTG
TGAAATTGTTATCCGCTCACAATTCCACACAACATACGAGCCGGAAGCATAAAGTGTAAAG
CCTGGGGTGCCTAATGAGTGAGCTAACTCACATTAATTGCGTTGCGCTCACTGCCCGCTTT
CCAGTCGGGAAACCTGTCGTGCCAGCGGATCCGCATCTCAATTAGTCAGCAACCATAGTC
CCGCCCCTAACTCCGCCCATCCCGCCCCTAACTCCGCCCAGTTCCGCCCATTCTCCGCCC
CATGGCTGACTAATTTTTTTTATTTATGCAGAGGCCGAGGCCGCCTCGGCCTCTGAGCTAT
TCCAGAAGTAGTGAGGAGGCTTTTTTGGAGGCCTAGGCTTTTGCAAAAAGCTAACTTGTTT
ATTGCAGCTTATAATGGTTACAAATAAAGCAATAGCATCACAAATTTCACAAATAAAGCATTT
TTTTCACTGCATTCTAGTTGTGGTTTGTCCAAACTCATCAATGTATCTTATCATGTCTGGATC
CGCTGCATTAATGAATCGGCCAACGCGCGGGGAGAGGCGGTTTGCGTATTGGGCGCTCTT
CCGCTTCCTCGCTCACTGACTCGCTGCGCTCGGTCGTTCGGCTGCGGCGAGCGGTATCA
GCTCACTCAAAGGCGGTAATACGGTTATCCACAGAATCAGGGGATAACGCAGGAAAGAAC
ATGTGAGCAAAAGGCCAGCAAAAGGCCAGGAACCGTAAAAAGGCCGCGTTGCTGGCGTTT
TTCCATAGGCTCCGCCCCCCTGACGAGCATCACAAAAATCGACGCTCAAGTCAGAGGTGG
CGAAACCCGACAGGACTATAAAGATACCAGGCGTTTCCCCCTGGAAGCTCCCTCGTGCGC
TCTCCTGTTCCGACCCTGCCGCTTACCGGATACCTGTCCGCCTTTCTCCCTTCGGGAAGC
GTGGCGCTTTCTCAATGCTCACGCTGTAGGTATCTCAGTTCGGTGTAGGTCGTTCGCTCCA
AGCTGGGCTGTGTGCACGAACCCCCCGTTCAGCCCGACCGCTGCGCCTTATCCGGTAACT
ATCGTCTTGAGTCCAACCCGGTAAGACACGACTTATCGCCACTGGCAGCAGCCACTGGTA
ACAGGATTAGCAGAGCGAGGTATGTAGGCGGTGCTACAGAGTTCTTGAAGTGGTGGCCTA
ACTACGGCTACACTAGAAGGACAGTATTTGGTATCTGCGCTCTGCTGAAGCCAGTTACCTT
CGGAAAAAGAGTTGGTAGCTCTTGATCCGGCAAACAAACCACCGCTGGTAGCGGTGGTTT
TTTTGTTTGCAAGCAGCAGATTACGCGCAGAAAAAAAGGATCTCAAGAAGATCCTTTGATCT
TTTCTACGGGGTCTGACGCTCAGTGGAACGAAAACTCACGTTAAGGGATTTTGGTCATGAG
ATTATCAAAAAGGATCTTCACCTAGATCCTTTTAAATTAAAAATGAAGTTTTAAATCAATCTAA
AGTATATATGAGTAAACTTGGTCTGACAGTTACCAATGCTTAATCAGTGAGGCACCTATCTC
AGCGATCTGTCTATTTCGTTCATCCATAGTTGCCTGACTCCCCGTCGTGTAGATAACTACGA
TACGGGAGGGCTTACCATCTGGCCCCAGTGCTGCAATGATACCGCGAGACCCACGCTCAC
CGGCTCCAGATTTATCAGCAATAAACCAGCCAGCCGGAAGGGCCGAGCGCAGAAGTGGTC
CTGCAACTTTATCCGCCTCCATCCAGTCTATTAATTGTTGCCGGGAAGCTAGAGTAAGTAG
TTCGCCAGTTAATAGTTTGCGCAACGTTGTTGCCATTGCTACAGGCATCGTGGTGTCACGC
TCGTCGTTTGGTATGGCTTCATTCAGCTCCGGTTCCCAACGATCAAGGCGAGTTACATGAT
CCCCCATGTTGTGCAAAAAAGCGGTTAGCTCCTTCGGTCCTCCGATCGTTGTCAGAAGTAA
GTTGGCCGCAGTGTTATCACTCATGGTTATGGCAGCACTGCATAATTCTCTTACTGTCATG
CCATCCGTAAGATGCTTTTCTGTGACTGGTGAGTACTCAACCAAGTCATTCTGAGAATAGT
GTATGCGGCGACCGAGTTGCTCTTGCCCGGCGTCAATACGGGATAATACCGCGCCACATA
GCAGAACTTTAAAAGTGCTCATCATTGGAAAACGTTCTTCGGGGCGAAAACTCTCAAGGAT
CTTACCGCTGTTGAGATCCAGTTCGATGTAACCCACTCGTGCACCCAACTGATCTTCAGCA

TCTTTTACTTTCACCAGCGTTTCTGGGTGAGCAAAAACAGGAAGGCAAAATGCCGCAAAAA
AGGGAATAAGGGCGACACGGAAATGTTGAATACTCATACTCTTCCTTTTTCAATATTATTGA
AGCATTTATCAGGGTTATTGTCTCATGAGCGGATACATATTTGAATGTATTTAGAAAAATAAA
CAAA

Green: left homology arm

Red: 5-nucleotide barcode

Dark Red: truncated CMV promoter

Light Blue: mKate open reading frame

Purple: PGK1 promoter

Blue: hygromycin resistance gene open reading frame

Orange: right homology arm

**Bray-Curtis and sequencing reads**

Assume that a PUF sample contains N barcode-indel reads, the average length of each read is L, and the error rate per base is e. Thus, the total number of mutations is $N * L * e$.

When $N * L * e << N$, each mutation most likely will occur within a different read. We further assume that the mutation does not result in a sequence identical to one of the original reads. Thus, for the $(N – N * L * e)$ non-mutated reads, they will appear in both the original and in the mutated samples. In contrast, for the $(N * L * e)$ mutated reads, they will only appear in the original sample.

Therefore, the Bray-Curtis value will be: $(N * L * e) / (N + N – N * L * e) = (L * e) / (2 – L * e)$.

Since $L * e << 1$, the Bray-Curtis value is $(L * e) / 2$, therefore the BC values are directly related to the read size L.

**Supplementary Tables**

**Supplementary Table 1. Primers used in this study**

**Supplementary Table 2. The list of individual barcodes and their frequencies for the pilot PUF.**

**Supplementary Table 3. The list of individual indels and their frequencies for the pilot PUF.**

**Supplementary Table 4. The PUF matrix for the pilot PUF.**

**Supplementary Table 5. The list of individual barcodes/indels and their frequencies for PUF1 samples.**

**Supplementary Table 6. The list of individual barcodes/indels and their frequencies for PUF2 samples.**

**Supplementary Table 7. The PUF matrices for PUF1 samples.**

**Supplementary Table 8. The PUF matrices for PUF2 samples.**

**Supplementary Table 9. The list of individual barcodes/indels and their frequencies for PUF3 samples.**

**Supplementary Table 10. The list of individual barcodes/indels and their frequencies for PUF4 samples.**

**Supplementary Table 11. The list of individual barcode-indel addresses and their frequencies for all PUF samples.**

**Supplementary Table 12. Total variation distances between PUF samples.**

**Supplementary Table 13. The Bray-Curtis dissimilarities between PUFs and their corresponding mutated samples.**

**Supplementary Table 14. The relative abundances of stably integrated barcodes in 6 replicates.**

**Supplementary Table 15. The Bray-Curtis dissimilarities between barcode replicates and their NGS sequencing replicates (denoted as r).**

**Supplementary Table 16. The Bray-Curtis dissimilarities between PUFs and their corresponding reshuffled samples.**
